# An Lrs14 family protein functions as a nucleoid-associated protein regulating cell cycle progression in Sulfolobales

**DOI:** 10.1101/2025.11.30.691466

**Authors:** Qi Gan, Haodun Li, Qihong Huang, Yunfeng Yang, Fei Sun, Xu Feng, Jinfeng Ni, Zhenfeng Zhang, Qunxin She, Yulong Shen

## Abstract

Archaea of the order Sulfolobales possess eukaryotic-like cell cycle. The chromatin organization in these archaea is supposed to rely on nucleoid-associated proteins (NAPs). Genomes of Sulfolobales species encode multiple members of the Lrs14 protein family, which have been postulated to function as NAPs. Here, we show that one of the Lrs14 family protein, named Sul14a, is a potential key player in dynamic chromatin organization in *Saccharolobus islandicus*. *Sul14a* is transcribed in a cell cycle-dependent manner and appears to be essential for cell viability. The protein exhibits DNA-binding activity preferring AT-rich sequences and can bridge DNA *in vitro.* ChIP-seq analysis revealed that Sul14a is enriched at AT-rich regions of the chromatin. Overexpression of Sul14a leads to cell cycle arrest at G2 phase. High-throughput chromosome conformation capture (Hi-C) and transcriptomic analyses indicate that Sul14a overexpression results in chromatin contact reduction and global transcriptional changes. In addition, we observed that the chromatin contact pattern changes markedly between 1h and 5h after release from G2 synchronization, corresponding to the lowest and highest levels of *sul14a* transcription, respectively, during the cell cycle. Our results establish that Sul14a is an NAP intimately associated with chromatin dynamics and global transcriptional changes during the cell cycle in Sulfolobales.

## Introduction

Archaea of the order Sulfolobales exhibit a eukaryotic-like cell cycle, with distinct phases comprising G1, S, G2, M, and D (cell division) phases. Species such as *Sulfolobus acidocaldarius*, *Saccharolobus islandicus*, and *Saccharolobus solfataricus* have been widely used as model organisms to investigate the biology and evolution of eukaryotic-like proteins and cellular processes, including DNA replication, chromatin compartmentalization, and cell division ^1–3^. Cell division in these archaea is controlled by the proteasome-mediated degradation of the cell division protein CdvB, whose timing is likely determined by the cyclically expressed ArsR-type repressor CCTF1 that regulates the expression of the proteasome-activating nucleotidase (PAN) ^3–6^. We have recently demonstrated that phosphorylation of the α subunit of the proteasome by the cyclically expressed kinase ePK2 inhibits the proteasome assembly and regulates the degradation of the cell division proteins CdvB, CdvB1, and CdvB2 in *Sa. islandicus* ^7^. In addition, recent work from our group and others has shown that the RHH domain-containing aCcr proteins regulates cell division by directly repressing several essential genes, including *cdvA* ^8–11^. However, many other aspects related to the cell cycle, including chromatin organization and gene regulation through chromatin dynamics are still unclear.

Chromosome organization is primarily shaped by chromatin proteins. Although histone homologs are absent in Sulfolobales archaea, these organisms possess several conserved, small, and abundant nucleoid-associated proteins (NAPs), including the canonical chromatin proteins Cren7, Sul7d, Sul10a, and Sul12a ^12–19^. The chromosomes of Sulfolobales archaea exhibit distinct structural features including chromosomal interaction domains (CIDs), A/B compartments, and loop structures ^2, 20, 21^. The Sulfolobales SMC family protein coalescin (ClsN) is enriched in transcriptionally inactive B compartments, where it likely contributes to chromatin organization by facilitating the bundling of CIDs ^2, 21^. However, how Sulfolobales archaea organize nascent DNA into such compartments through chromatin proteins and dynamically regulate transcription in an orderly manner remains poorly understood.

The Lrs14 protein was first identified in *Sa. solfataricus* (formerly *Sulfolobus solfataricus*) ^22^. It contains a helix-turn-helix motif and belongs to the leucine-responsive regulatory protein (Lrp-AsnC) family. Lrs14 binds multiple sites within its own promoter, and its transcript accumulates during the late growth stages of *Sa. solfataricus* ^22^. Since then, homologs and orthologs of Lrs14 have been identified and assigned different names, including Sta1 and Smj12 in *Sa. solfataricus* ^23–25^, AbfR1 and AbfR2 in *S. acidocaldarius* ^26, 27^. Now it is known that Lrs14-like proteins are widespread in members of the Sulfolobales ^28^. Across the genomes of the representative Sulfolobales genera, four to nine Lrs14 orthologs are predicted to be encoded. Although initially classified as transcription factors, these proteins are actually unresponsive to a specific ligand and lack a canonical ligand-binding domain ^28^. The Lrs14 family member AbfR1, which has a specific regulatory role in biofilm formation and cell motility in *S. acidocaldarius*, has been proposed to act as a global DNA-binding protein that combines pleiotropic transcriptional regulation with a chromatin-structuring role (Peeters et al., 2015). Based on a comprehensive analysis of available data on Lrs14 family proteins, including their high abundance, structural similarity to known chromatin proteins (Sso10a, Sul12a, and TrmBL2), and their non-specific DNA-binding activity that modulates DNA architecture, it has been postulated that the Lrs14 family members act as nucleoid-associated proteins in Sulfolobales. They are proposed to fulfill dual roles in DNA structuring and global transcriptional regulation, particularly under stress conditions ^28^. However, the mechanisms by which these proteins organize chromatin and regulate gene expression in response to cellular signals and environmental cues remain unclear.

In this study, we characterized one member of the Lrs14 proteins, named Sul14a, from *Sa. islandicus,* both *in vitro* and *in vivo*. The *sul14a* gene is transcribed in a cell cycle-dependent manner and appears to be essential for cell viability. The protein exhibits DNA binding activity with a preference for AT-rich sequences and displays DNA-bridging activity *in vitro.* Genome-wide ChIP-Seq analysis further revealed its enrichment at AT-rich regions *in vivo*. Integrative Hi-C, ChIP-seq, and transcriptomic analyses demonstrate that Sul14a is closely associated with chromatin dynamics during the cell cycle, suggesting that it plays an essential role in coordinating orderly cell cycle progression in Sulfolobales archaea.

## Results

### *Sul14a* is transcribed in a cell cycle-dependent pattern and is essential for cell viability

The genome of *Sa. islandicus* REY15A encodes six Lrs14-like proteins ^29^ (Table S1). The activities, characterized mostly *in vitro*, and structural information of several homologs have been reported in the closely related species ^28^. One of our main interests is the cell cycle regulation mechanism in Sulfolobales archaea. A genome-wide microarray analysis showed that the *lrs14* homolog in *S. acidocaldarius*, *saci_0102*, is transcribed in a cell cycle-dependent manner ^30^. The homolog of Saci_0102 in *Sa. solfataricus* (SSO0048) was pulled down using probes representing promoter regions of rudivirus SIRV1 genes and the *radA*-paralog gene (*sso0777*) as baits, and was named as Sta1 (Sulfolobus transcription activator 1) for its ability to activate viral gene transcription ^23, 24^. To elucidate the function and mechanism of Lrs14 family proteins in the cell cycle of Sulfolobales, we focus on the Saci_0102 homolog in the model archaeon *Sa. islandicus* REY15A. This protein, hereafter referred to as Sul14a, is encoded by *sire_1948* (390 bp, 129 aa). Multiple sequence alignment of representative sequences from 28 Sulfolobales species show that these sequences are highly conserved, with an average pairwise identity of 76.5%, especially at the core DNA-binding domain (Fig. S1a). The phylogenetic tree of the Lrs14 homologs recapitulated the species tree of Sulfolobales (Fig. S1b), implying a conserved and important role of these proteins in Sulfolobales archaea.

Our previous transcriptomic data of synchronized *Sa. islandicus* E233S cells showed that *sul14a* is transcribed in a cell cycle-dependent manner ^8^. To validate this and to prepare samples for further analysis, we performed synchronization of the cells as before ^8^ (Fig. 1a). After release from G2 synchronization, the cell cycle proceeded through M-phase to D-phase, as observed by flow cytometry. Cells containing a single-copy chromosome (1C) gradually dominated the population by 3-4 h, completing one cycle upon re-entry into G2-phase (Fig. 1a,b). As shown in Fig. 1c, the transcription of *sul14a* increased from 1 h to 5 h, peaked at 5 h, and declined at 6 h. To know whether Sul14a protein levels followed a similar pattern, we performed Western blotting analysis. Interestingly, the protein levels remained largely constant throughout the cell cycle (Fig. 1d), in sharp contrast to the cyclic transcriptiona profile.

**Fig. 1.**
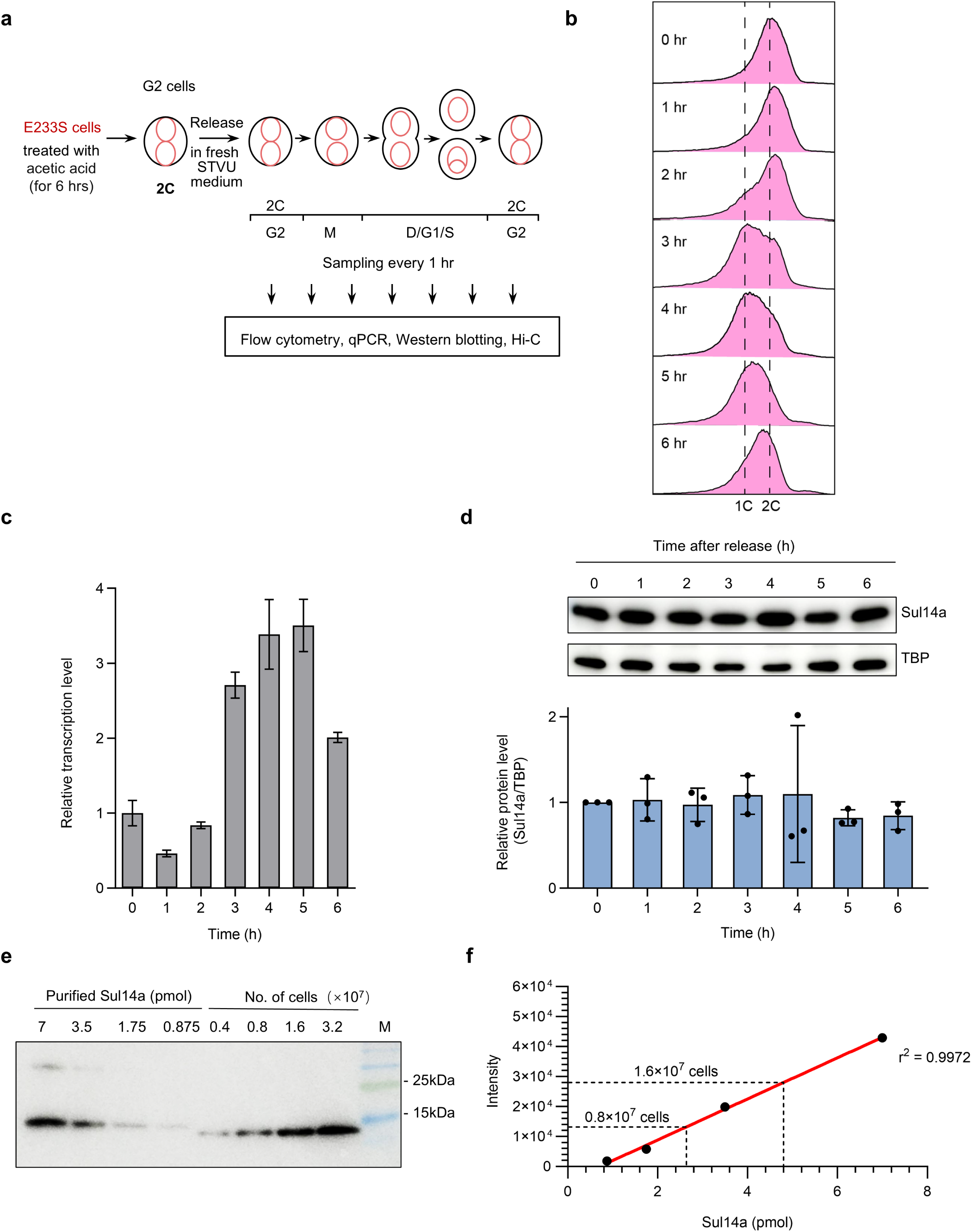
*Sul14a* exhibits a cyclic transcription pattern in *Sa. islandicus* REY15A. **a,** Schematic of the experimental workflow for cell cycle synchronization and time-point sampling of E233S cells. **b**, Flow cytometry profiles of E233S cells after synchronization. Cells (200,000 per sample) were collected for the analysis. **c,** Relative transcription levels of *sul14a* during one cell cycle measured by qRT-PCR. The gene *tbp* was used as a control and set as 1.0. The values were based on three technical repeats. **d,** Western blotting analysis of Sul14a levels during a cell cycle using anti-Sul14a antibody. TBP (TATA-box binding protein) was used as a control. A representative image was shown. The panel below shows quantification of the results by ImageJ. The values were measured based on at least three technical repeats. **e,** Estimate of the amounts of Sul14a protein in the cells by Western blotting using anti-Sul14a antibody. Recombinant Sul14a and cell extracts of E233S were subject to the analysis. **f,** Calibration curves of band intensity and the amounts of recombinant Sul14a in (**e**). The quantitation was performed using ImageJ.

To assess the functional importance of *sul14a*, we attempted to generate a gene knockout mutant. Despite multiple attempts, no viable deletion mutant was obtained, indicating that *sul14a* is likely essential for cell viability in *Sa. islandicus*. This finding is consistent with a previous genome-wide transposon mutagenesis study in *Sa. islandicus* ^31^. We then quantified the intracellular abundance of Sul14a by Western blotting, using recombinant protein as a calibration standard. Each cell contains approximately 1.8×10⁵ Sul14a as monomers (Fig. 1e, f), corresponding to a mass ratio of 1.5:1 relative to genomic DNA. This suggests that one Sul14a dimer would theoretically cover 28 bp of DNA, assuming complete chromosomal binding. For comparison, the known chromatin protein Cren7 was reported to have a mass ratio of 0.8:1 in *Sa. solfataricus*, constituting approximately 1% of total cellular protein ^12^. These results support the notion that Sul14a is an essential NAP, required for cell cycle progression and cell survival.

### Sul14a preferentially binds AT-rich DNA and exhibits DNA bridging activity *in vitro*

To examine the DNA-binding properties of Sul14a, the protein was expressed and purified from *E. coli,* and analyzed by size exclusion chromatography (SEC) and *in vitro* chemical cross-linking (Fig. 2a and Fig. S2a,b). The results confirmed that Sul14a forms a dimer in solution. Sul14a primarily forms dimer with different protein amounts (3 or 6 μg) in the reactions, while more higher-order oligomer bands were observed with elevated more protein (6 μg) in the reactions (Fig. S2b). The presence of DNA further promoted the formation of higher-order oligomers (Fig. S2c), indicating that DNA binding facilitates Sul14a oligomerization. EMSA revealed that Sul14a can bind both linear and circular plasmid dsDNA (Fig. 2b and Fig. S2d). Notably, Sul14a exhibits a preference for AT-rich DNA, as demonstrated using a substrate with 100% AT content (Fig. 2c). Sul14a bound the AT-rich substrate with a dissociation constant (Kd) of 0.375 ± 0.047 µM, whereas its binding to a GC-rich dsDNA (100% GC) was substantially weaker and did not reach saturation even at 9.6 µM protein (Fig. 2d). These results indicate that Sul14a binds DNA in a non-sequence-specific manner and prefers AT-rich DNA *in vitro*. Overall, these findings demonstrate that Sul14a shares the characteristic DNA-binding properties of the Lrs14 family proteins.

**Fig. 2.**
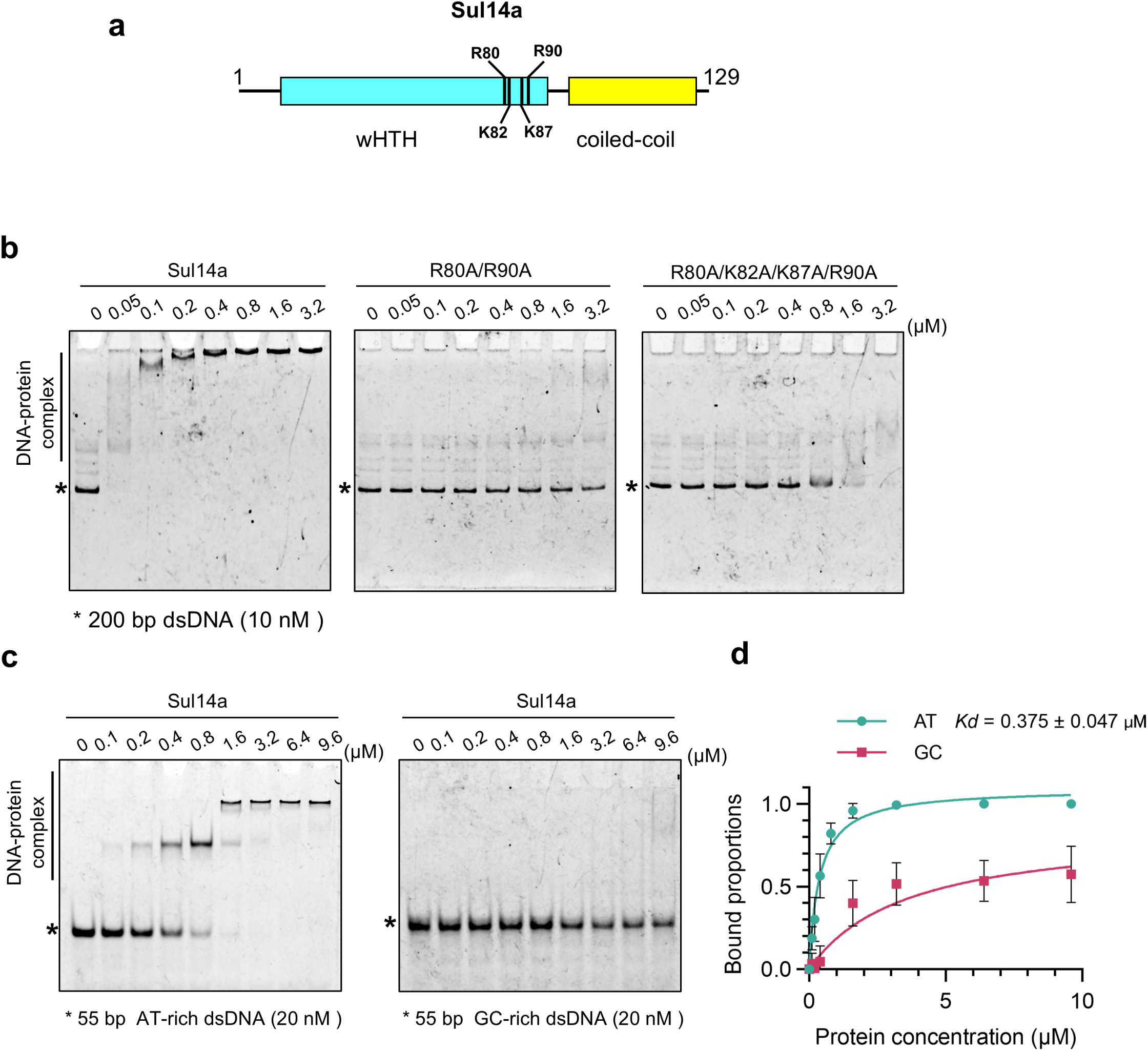
Sul14a binds DNA non-specifically with AT preference *in vitro*. **a,** Domain organization of Sul14a. Sul14a is a small protein of 129 amino acids that contains an N-terminal wHTH domain and a C-terminal coiled-coil domain. The black bars indicate the identified DNA binding sites. **b,** Analysis of the DNA binding capacity of Sul14a and its mutants by EMSA. The assay was performed with 10 nm DNA and increasing concentrations of Sul14a (0, 0.05, 0.1,0.2, 0.4, 0.8, 1.6, and 3.2 µM). **c,** Sul14a exhibits differential DNA binding capacity for AT-rich and GC-rich sequences. Increasing concentrations of Sul14a (0, 0.1, 0.2, 0.4, 0.8, 1.6, 3.2, 6.4, and 9.6 µM) were incubated with the substrate DNA at a fixed final concentration of 20 nM. **d,** Quantitative analysis of the results in (**c**). The quantitation was performed using ImageJ, with the fitted curve and binding constant derived from the Michaelis-Menten equation. At least two technical repeats were performed for each set of the EMSA assay.

To clarify the structural basis of its biochemical properties, we predicted the 3D structure of Sul14a using AlphaFold Server ^32^. Each monomer consists of an N-terminal winged helix-turn-helix (wHTH) domain (α-helices H1–H4 and two β-sheets) responsible for DNA binding, and a C-terminal coiled-coil domain (α-helix H5) (Fig. 2a and S3a). Homology modeling using *S. acidocaldarius* AbfR2 (30.8% identity, 51.9% sequence similarity) (PDB:6CMV) (Fig. S3c) and *S. tokodaii* Sto12a (34.4% identity, 50.4% sequence similarity) (PDB:2D1H) ^33^ suggested a dimeric assembly of Sul14a (Fig. S3b,c). We identified highly conserved residues R80, K82, K87, and R90 at the positively charged loop (β3-β4), predicted to be involved in DNA binding (Fig. S1a and Fig. S3d) and subsequently expressed and purified various alanine-substitution mutants (Fig. S4a). EMSA results showed that only the R80A/R90A and R80A/K82A/K87A/R90A mutants exhibited substantially reduced DNA-binding activity compared to the wild-type protein (Fig. 2b). In contrast, single mutants (R80A, K82A, K87A, R90A) and other double mutants (R80A/K82A, K82A/K87A, K87A/R90A) largely retained their DNA-binding capacity (Fig. S4b,c). These results demonstrate that R80 and R90 are critical for DNA binding and function cooperatively at the binding interface, presumably by stabilizing the protein–DNA complex. Structural prediction suggests that R90 could form multiple hydrogen bonds with the DNA phosphate backbone within the minor groove, with R80 synergistically supporting this interaction with adjacent basic residues (Fig. S3d).

Some chromatin proteins in bacteria have DNA bridging activity to facilitate DNA packaging ^34, 35^. We analyzed dsDNA bridging ability following a fluorescent-labeled DNA pull-down assay ^36, 37^ (Fig. 3a). Briefly, magnetic streptavidin beads were incubated with a biotin-labeled bait DNA, then Sul14a and FAM-labeled (500 bp) prey DNA substrate were added. If Sul14a is able to bridge dsDNA, the prey DNA substrate will be eluted and detected with the bait DNA after wash. The results showed that Sul14a is able to bridge diverse DNA substrates, including a 500-bp linear dsDNA, and a 4.3-kb linearized, supercoiled, and relaxed plasmid DNA (Fig. 3b,c). This activity relies on the DNA binding activity, because the DNA-binding deficient mutants R80A/R90A and R80A/K82A/K87A/R90A completely lost the bridging activity (Fig. 3b). The bridging activity of Sul14a facilitates its function as a potent chromatin organizer.

**Fig. 3.**
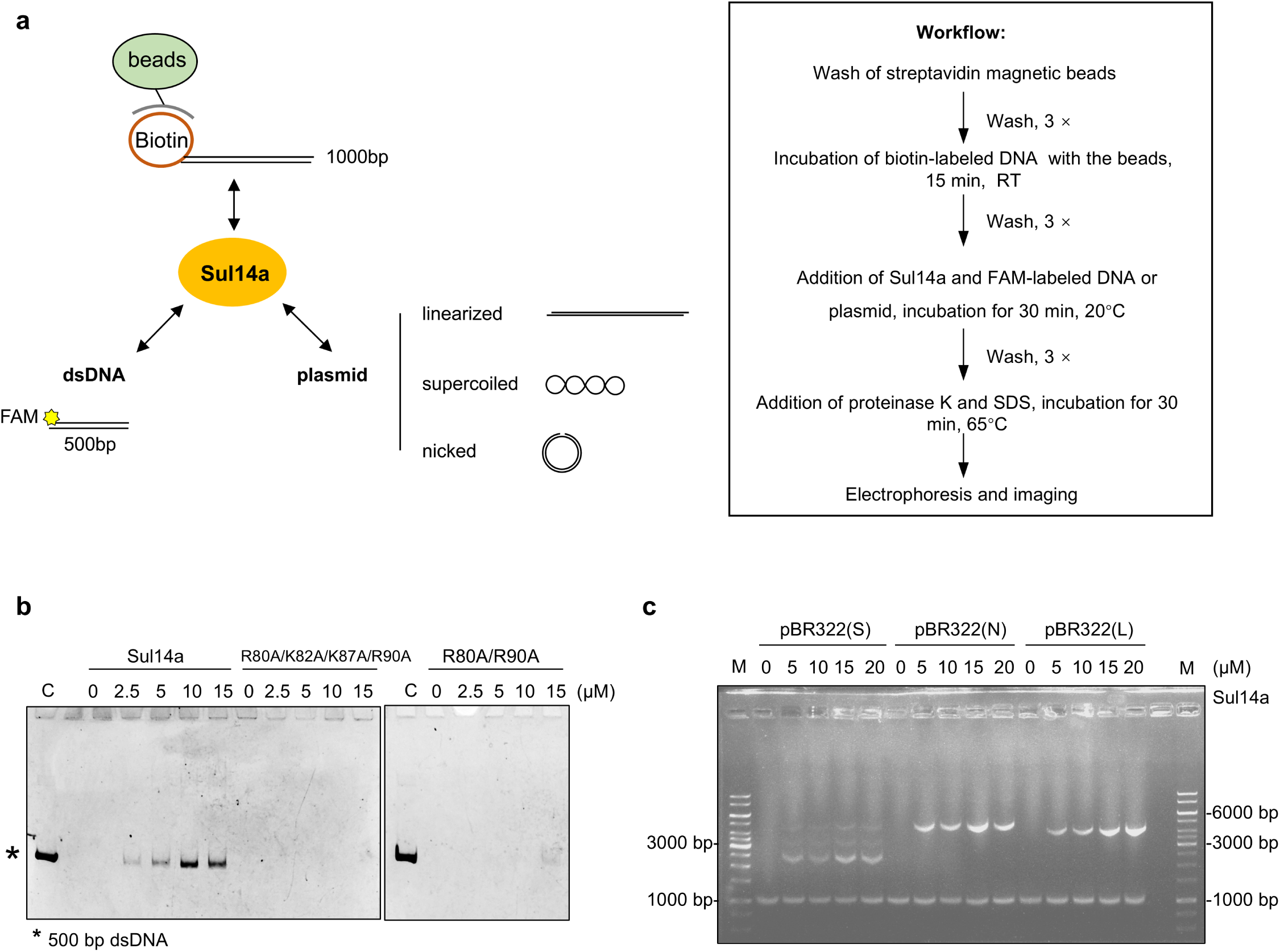
Sul14a bridges DNA. **a,** Schematic and workflow showing the principle and procedure for the DNA bridging assay. Magnetic streptavidin beads were incubated with the biotin-labeled bait 1,000 bp DNA (200 ng per reaction) before Sul14a was added. FAM-labeled (500 bp) double-stranded linear or plasmid DNA with three topological states were used as the bridging substrates. **b,** Bridging ability of Sul14a and its DNA-binding deficient mutants using 500 bp linear dsDNA. C, control. Prey DNA (15 nM) was used for each reaction. **c,** Bridging ability of Sul14a using linearized (L), supercoiled (S), and nicked (N) plasmid DNA. Each reaction was performed with at least two technical replicates.

### Overexpression of Sul14a impairs cell proliferation and cell cycle progression

Due to unsuccessful attempts to generate *sul14a* knockout or knockdown strains based on the endogenous CRISPR-Cas-based genetic system ^38, 39^, we constructed Sul14a overexpression strains in *Sa. islandicus* E233S to investigate its function. Three types of strains were constructed: the wild-type (pSeSD-Sul14a), C-terminal His-tagged wild-type (pSeSD-Sul14a-C-His), and C-terminal His-tagged DNA-binding deficient mutants with single/double/quadruple substitutions (R80A, K82A, K87A, R90A and their combinations). Cells were cultured in STV (low-induction) and ATV (induction) media, followed by growth curve and flow cytometry analyses (this section), as well as ChIP-seq, RNA-seq, and Hi-C experiments using both unsynchronized and G2–phase–synchronized cells (see below) (Fig. 4a). In ATV medium, the *sul14a* transcript levels of pSeSD-Sul14a cells were elevated approximately 25- to 40-fold compared with control cells carrying the empty vector pSeSD (Fig. S5a,b), whereas the protein levels increased by only about threefold after induction (Fig. S5c,d). The expression of Sul14a proteins in all strains were confirmed by Western blotting (Fig. S5e). All strains exhibited normal growth in STV medium similar to the control (pSeSD) (Fig. 4b and Fig. S5f). Upon arabinose induction, however, growth of strains carrying pSeSD-Sul14a, pSeSD-Sul14a-C-His, pSeSD-Sul14a-K82A-C-His, pSeSD-Sul14a-K87A-C-His) was severely inhibited. In contrast, cells expressing the DNA-binding deficient mutants R80A/R90A and R80A/K82A/K87A/R90A displayed growth comparable to that of the control cells (Fig. 4b and Fig. S5g). These results indicate that overexpression of Sul14a strongly impairs cell growth in a manner dependent on its DNA-binding/bridging activity. Notably, the addition of a C-terminal His-tag for ectopic expression does not compromise its function.

**Fig. 4.**
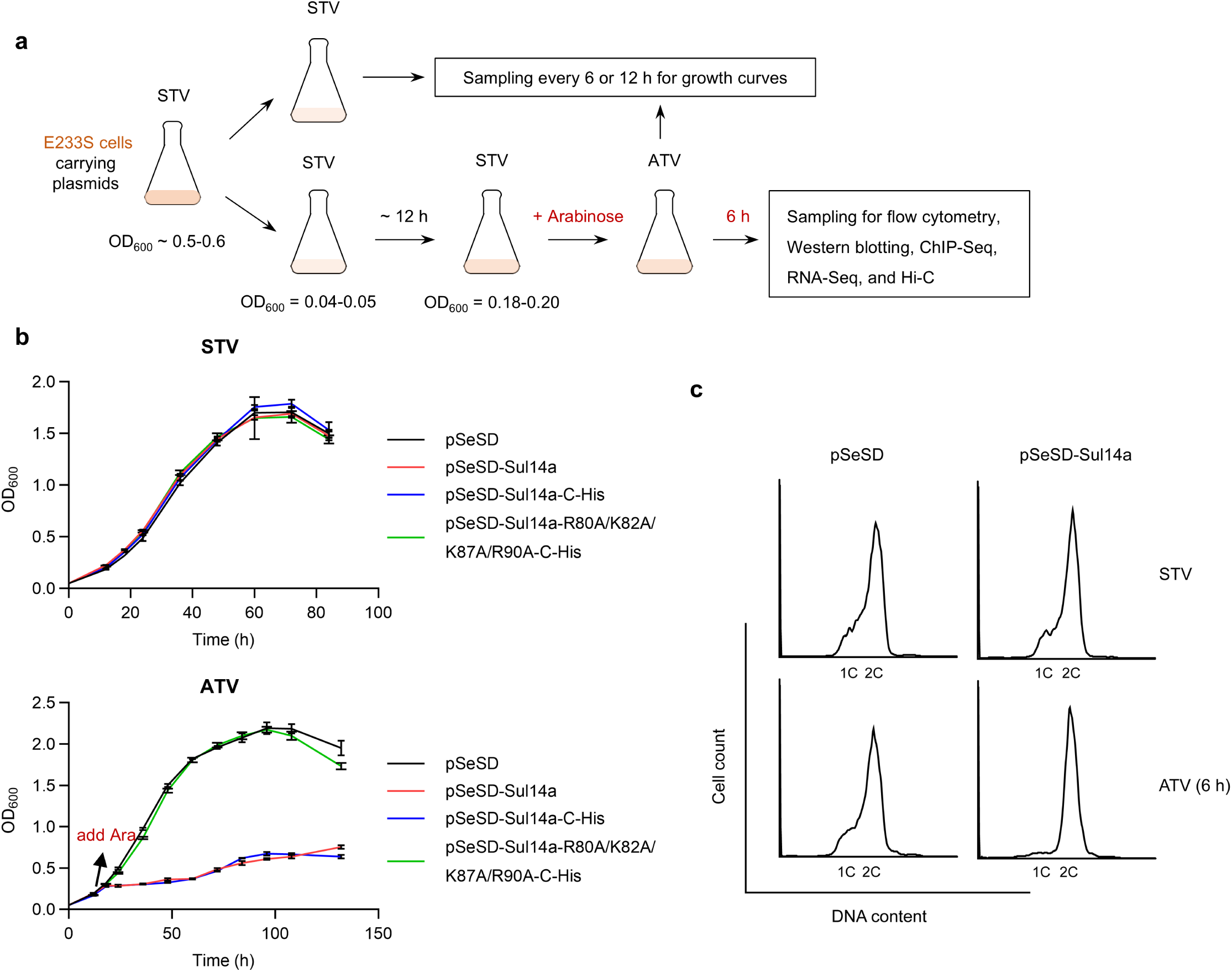
Sul14a overexpression inhibits cell growth of *Sa. islandicus* significantly. **a**, Schematic of the experimental workflow and sampling strategy for the analysis of the Sul14a overexpression strains. **b**, Growth curves of Sul14a and its DNA-binding deficient mutant overexpression strains cultivated in STV (upper panel) or ATV (lower panel) medium. **c**, Flow cytometry profiles of the Sul14a overexpression strains (unsynchronized) cultivated in STV or ATV medium after 6 h induction with arabinose. Cells containing one and two chromosomes (1C and 2C) are indicated. Cells carrying pSeSD were used as the control.

Flow cytometry analysis of unsynchronized cells showed that in STV medium, the DNA content profile of the pSeSD-Sul14a cells was indistinguishable from that of the control (pSeSD) at 6 h. However, when cultured in ATV medium, the 1C population was completely depleted, while the 2C population retained (Fig. 4c). These results indicate that overexpression of Sul14a severely disrupts normal cell cycle progression. To further assess the impact of Sul14a overexpression on cell cycle progression, cells were synchronized to G2 phase using acetic acid treatment. Overexpression was induced by adding arabinose 2 hours before release, and samples were collected hourly for flow cytometry analysis (Fig. 5a). As expected, Sul14a overexpression resulted in a blockade of chromosome segregation and/or cell division, with cells remaining arrested in G2 phase throughout the observation period (Fig. 5b). The observed G2 arrest suggests that Sul14a overexpression disrupts chromatin normal dynamics required for G2/M(D) transition, a process that probably needs to be tightly controlled as in eukaryotes.

**Fig. 5.**
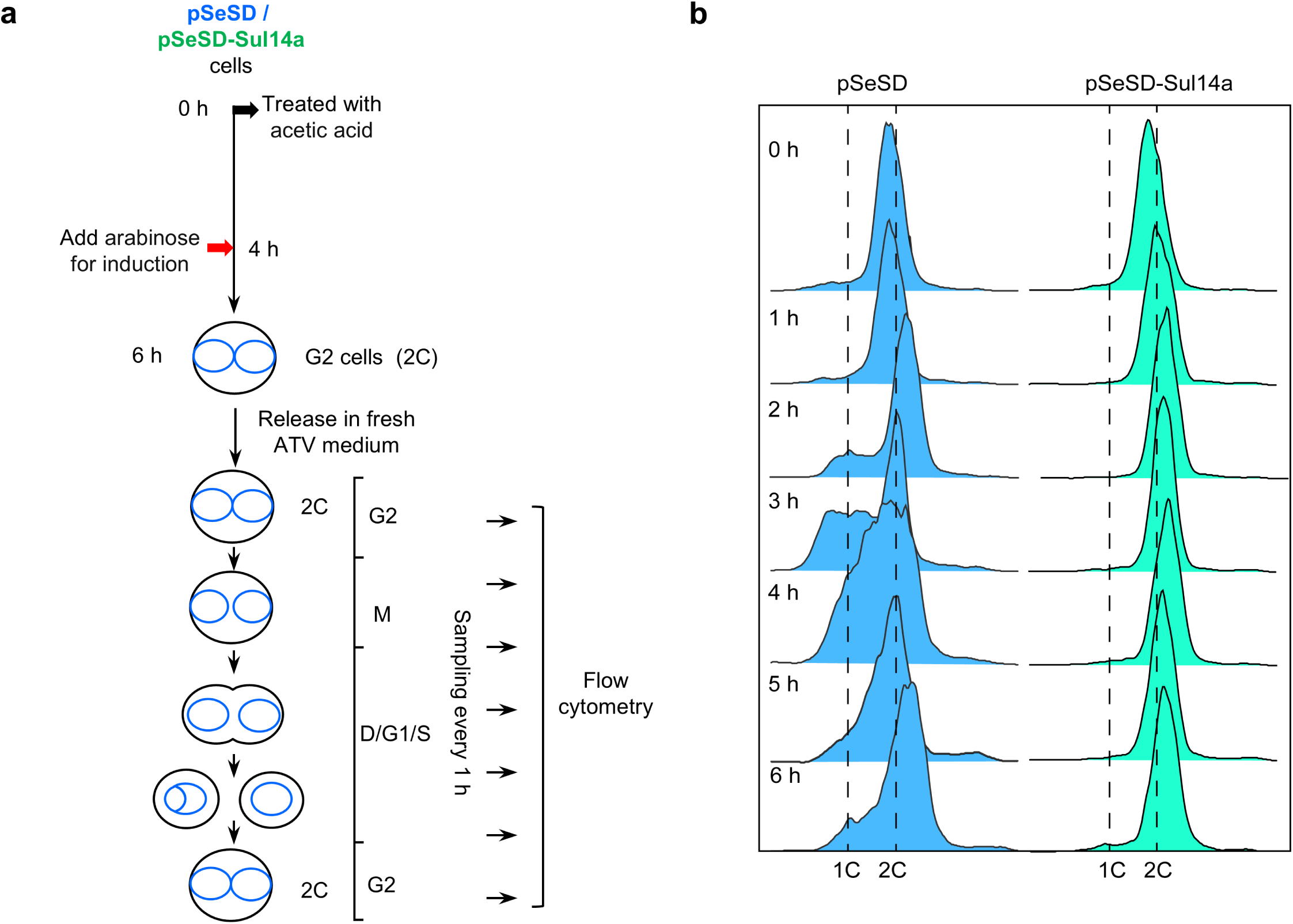
Sul14a overexpression arrests the cell cycle of *Sa. islandicus* at the G2 phase. **a,** Schematic of the experimental design and sampling strategy for cell cycle synchronization of the Sul14a overexpression strain. Arabinose was added 2 h before release for the induction of Sul14a. **b,** Flow cytometry profiles of Sul14a overexpression and the control strains after release. Cells containing one and two chromosomes (1C and 2C) are indicated. Each set of experiments was conducted at least twice.

### Sul14a preferentially binds AT-rich regions *in vivo*

To investigate the mechanism by which Sul14a overexpression affects cell cycle progression, we performed chromatin fractionation analysis on E233S cells. The results revealed that Sul14a is predominantly localized in the chromatin fraction during both exponential and stationary growth phases, with only a minor portion detected in the soluble fraction (Fig. S6a). Consistently, DNase I treatment markedly reduced the amount of Sul14a in the chromatin fraction (Fig. S6a). We next performed ChIP-Seq analysis on cells carrying pSeSD and pSeSD-Sul14a (Fig. 6a,c). The results showed that Sul14a is broadly enriched across the chromosome. A total of 780 and 802 high-confidence Sul14a enrichment peaks (FDR < 0.05) were identified in the control and overexpression samples, with average widths of 968 bp and 1,069 bp, respectively (Supplementary Data 1). Genome-wide profiling revealed that >80% of these peaks are located in promoter-proximal regions (Fig. S6b). Given that Sul14a preferentially binds AT-rich sequences *in vitro*, we next analyzed the correlation between genomic AT content and Sul14a enrichment. Across the genome, Sul14a enrichment showed a strong positive correlation with AT content in pSeSD cells (Spearman *r* = 0.66, *P* < 0.0001), which became even stronger in the overexpression cells (Spearman *r* = 0.83, *P* < 0.0001) (Fig. 6b,d and S6c). To further characterize the binding patterns, we compared Sul14a ChIP-seq enrichment among three genomic regions, promoter (TSS-proximal, 65.2% AT), gene bodies (mid-50%, 63.8% AT), and convergent intergenic regions (n=263, 69.8% AT), using 1.0-kb flanking windows (Fig. 6e–g). In both control and Sul14a-overexpressing cells, enrichment was consistently higher at the TSS than within gene bodies. Notably, the convergent intergenic regions, which have the highest average AT content, exhibited the strongest and most uniform Sul14a binding. Upon Sul14a overexpression, enrichment at the convergent intergenic regions decreased, whereas promoter-proximal binding increased markedly. These results confirm that Sul14a preferentially binds AT-rich sequences *in vivo*. The shift toward promoter occupancy at elevated protein levels suggests that Sul14a modulates gene expression by selectively binding promoter-proximal regions when its cellular abundance increases. Furthermore, the pronounced enrichment of Sul14a in AT-rich convergent intergenic regions supports a role in higher-order chromatin organization, potentially through mediating long-range DNA interactions or regional compaction. We also compared the distribution of Sul14a ChIP-seq enrichment between the A and B compartments. As shown in Fig. S6d,e, the A compartment showed slightly higher Sul14a enrichment than the B compartment in both pSeSD and pSeSD-Sul14a cells.

**Fig. 6.**
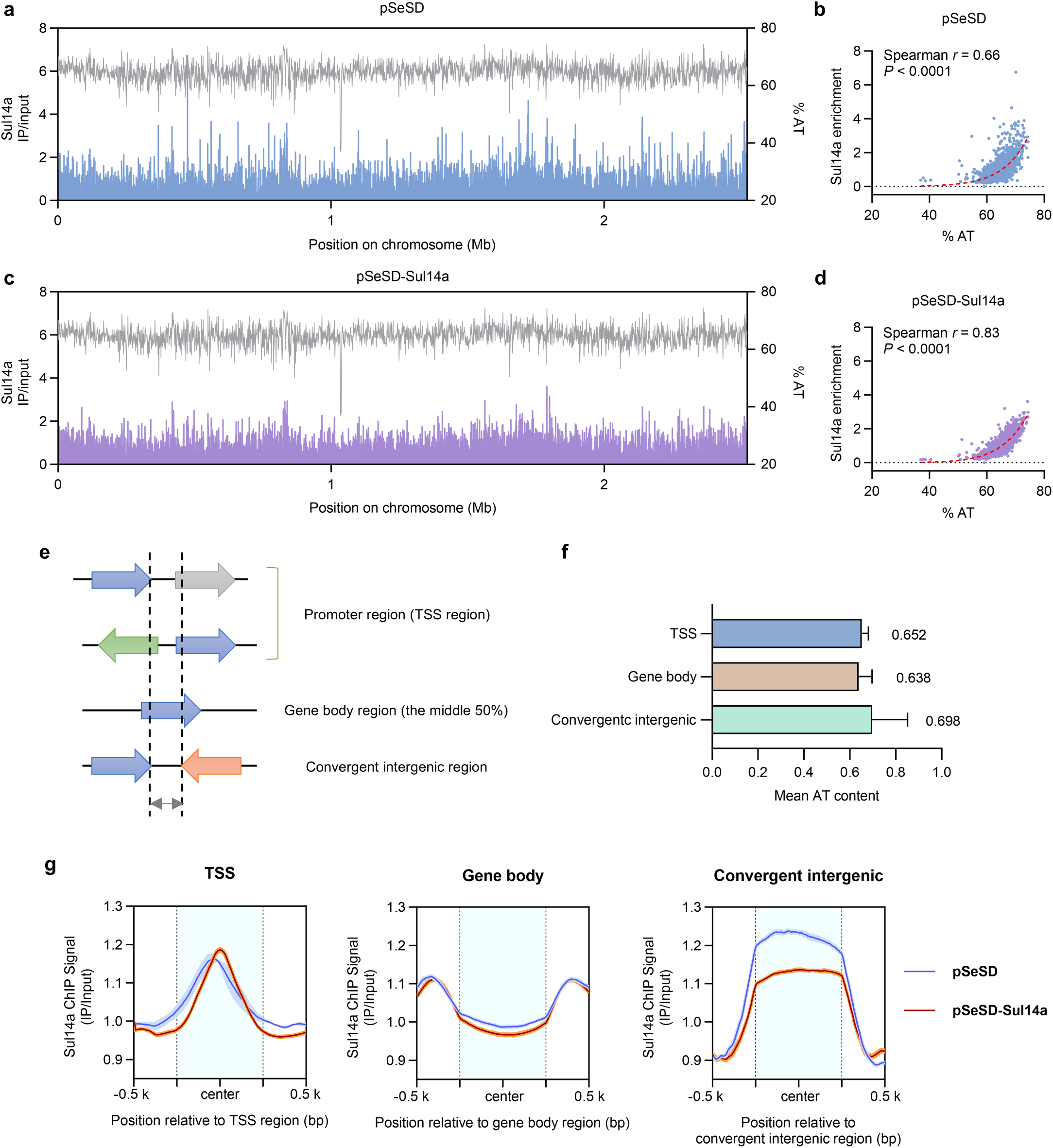
Genome-wide ChIP-seq profiling demonstrates the enrichment of Sul14a at AT-rich regions *in vivo.* **a,** ChIP-Seq analysis of Sul14a enrichment on *Sa. islandicus* REY15A chromosome in the control (pSeSD) strain. All ChIP data are normalized to input DNA. Genome coordinates are shown on the x-axis. The AT content coordinates are shown on the right y-axis. Two biological replicates were used for the analysis. The data were analyzed at 1-kb resolution. **b,** Correlation between the Sul14a enrichment in (a) and the AT content (%) of *Sa. islandicus* REY15A genome. Spearman correlations (*r*) were calculated and a Gaussian fitting was performed to obtain the curve (red dashed line). The data were analyzed at 1-kb resolution. **c,** ChIP-Seq analysis of Sul14a enrichment on *Sa. islandicus* REY15A chromosome in the Sul14a overexpression (pSeSD-Sul14a) strain. **d,** Correlation between the Sul14a enrichment in (c) and the AT content (%) of *Sa. islandicus* REY15A genome. **e,** Schematic diagrams of three genomic regions: the promoter region (TSS region), gene body region, and the convergent intergenic region. The middle 50% of the gene body region was selected for analysis. **f,** The mean AT content of the three regions. **g,** Sul14a ChIP signal enrichment in the three genomic regions indicated in (e) (1-kb windows). Plots show average signal intensity (y-axis) versus relative genomic position (x-axis, bp). The line represents the mean (IP/input with SES scaling). The shaded area represents the 95% confidence interval.

### Overexpression of Sul14a results in profound chromatin compartmentalization changes

To gain more insights into the role of Sul14a as a chromatin-associated protein, we performed Hi-C analysis on cells carrying pSeSD and pSeSD-Sul14a. The Hi-C contact matrix of the control (pSeSD) cells exhibited the characteristic plaid-like pattern as previously described for *Sa. islandicus* ^2^ (Fig. 7a). Strikingly, in the Sul14a-overexpressing strain, the boundaries of the plaid pattern became less distinct, and chromatin interactions along the primary diagonal were noticeably weakened (Fig. 7a). To assess the effects on chromatin compartmentalization, we calculated Pearson correlation coefficients based on the Hi-C contact maps and derived the compartment indices through principal component analysis (PCA). The first principal component (PC1) was used to assign genomic regions to the transcriptionally active A compartment (positive PC1 values) and the inactive B compartment (negative PC1 values) (Fig. 7b, lower panels). The Pearson correlation matrices showed that the composition of A and B compartments became blurred upon Sul14a overexpression, and this was accompanied by reciprocal shift between A and B compartments (Fig. 7b). We generated a Hi-C contact matrix showing the ratio of interaction frequencies between Sul14a-overexpressing and the control cells. As shown in Fig. 7c, Sul14a overexpression induced pronounced changes in chromatin compartmentalization. Notably, Sul14a overexpression appeared to enhance inter-domain (A-B) interactions but weaken intra-domain (A-A and B-B) interactions. Consistently, the contact frequency distributions between the two strains showed reduced close- to mid-chromatin interactions but a marked increase in long-range interactions in Sul14a-overexpressing cells (Fig. 7d). Together, these findings indicate that Sul14a overexpression profoundly disrupts higher-order chromosome architecture, leading to weakened compartmentalization and increased long-range chromatin contacts.

**Fig. 7.**
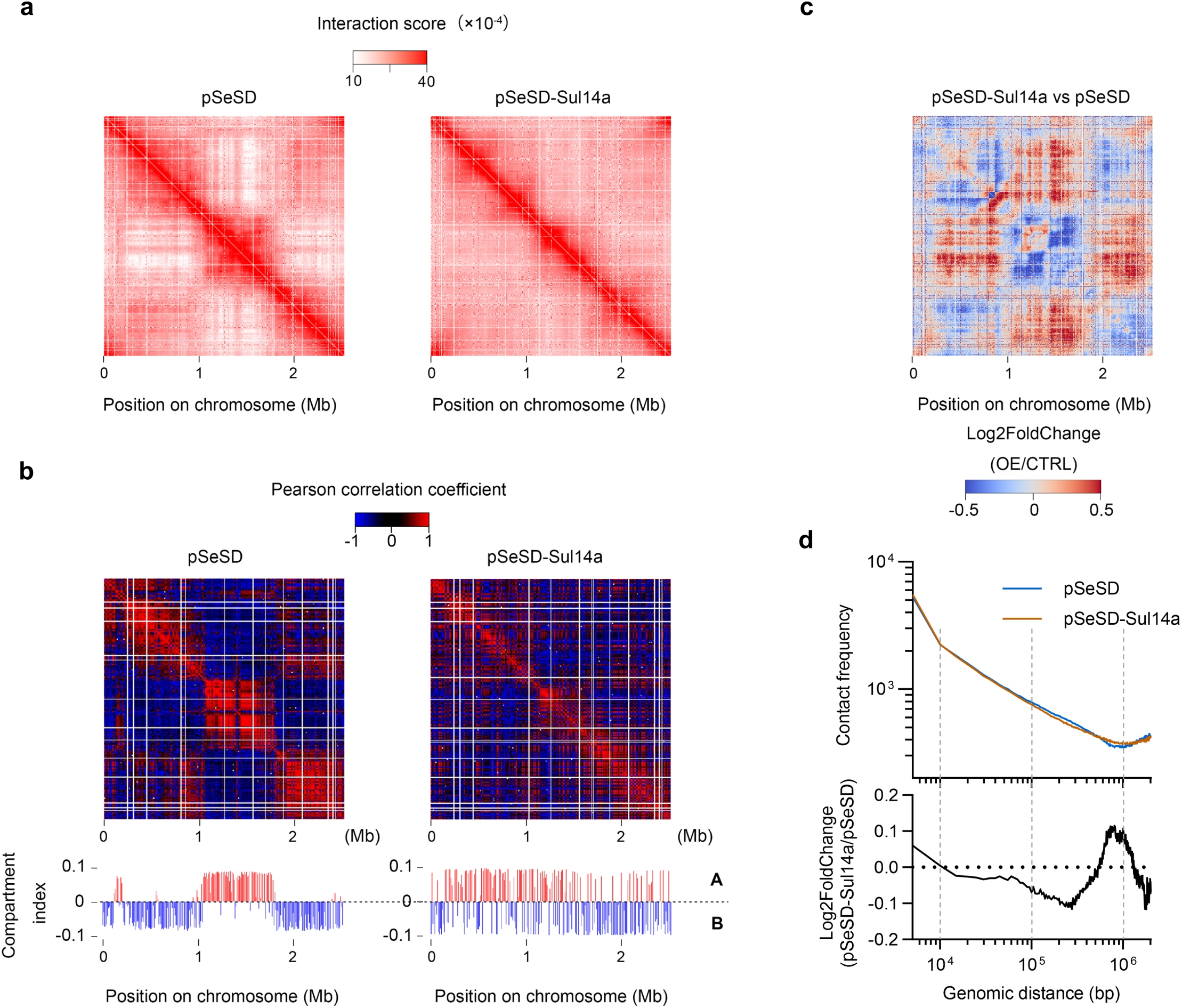
Overexpression of Sul14a resulted in profound chromatin compartmentalization changes. **a**, Hi-C contact matrices of cells carrying pSeSD and pSeSD-Sul14a at 5-kb resolution. **b**, Pearson correlation matrices of Hi-C data at 10-kb bin. The bottom panels show the compartment index matrices generated PCA based on the Pearson correlation data. The data were combined from two biological replicates. The A and B compartments are showed in red and blue, respectively. **c**, Matrices of the ratios of Hi-C interaction scores between pSeSD and pSeSD-Sul14a strains in (a). **d,** Long-range chromosomal contacts increased in the control and the Sul14a overexpression cells. Upper panels, the contact frequencies in pSeSD (blue) and pSeSD-Sul14a (orange) cells are plotted as a function of genomic distances (log scale; biological replicates combined). Bottom panels, Log2 fold changes in contact frequencies between comparative groups are shown against log-scaled genomic distances. The data were analysis at 5-kb bin.

### Analysis of chromatin compartmentalization in G2-synchronized cells during peak and trough transcription of *sul14a*

We observed that the *sul14a* gene exhibits a clear cell cycle-dependent transcriptional pattern, with the lowest expression at 1 h (trough) and the highest at 5 h (peak) after release from G2 synchronization (Fig. 1a-c). Although Sul14a protein levels remained relatively constant (Fig. 1d), we reasoned that the chromatin-bound fraction of Sul14a might fluctuate during the cell cycle, possibly regulated by post-translational modification. Based on this, we hypothesized that chromatin architecture could differ between the trough and peak phases of *sul14a* transcription. To test this, we performed Hi-C analysis using exponentially growing asynchronous E233S cells (CK) and G2-synchronized cells collected at 1 h (SC-1h) and 5 h (SC-5h) after release. The contact matrices of E233S cells displayed a clear plaid pattern, whereas the pattern boundaries appeared less distinct in SC-1h and SC-5h cells (Fig. 8a). All three samples exhibited similar A/B compartmentalization profiles, with the control (asynchronous E233S) showing the most pronounced pattern. Notably, chromatin compartment boundaries near the main diagonal appeared relatively diffuse during the trough (SC-1h) but became increasingly distinct at the peak (SC-5h) transcription, with the overall compartmentalization pattern resembling that of the control (Fig. 8a,b). These observations suggest that chromatin undergoes subtle yet dynamic reorganization during the cell cycle.

**Fig. 8.**
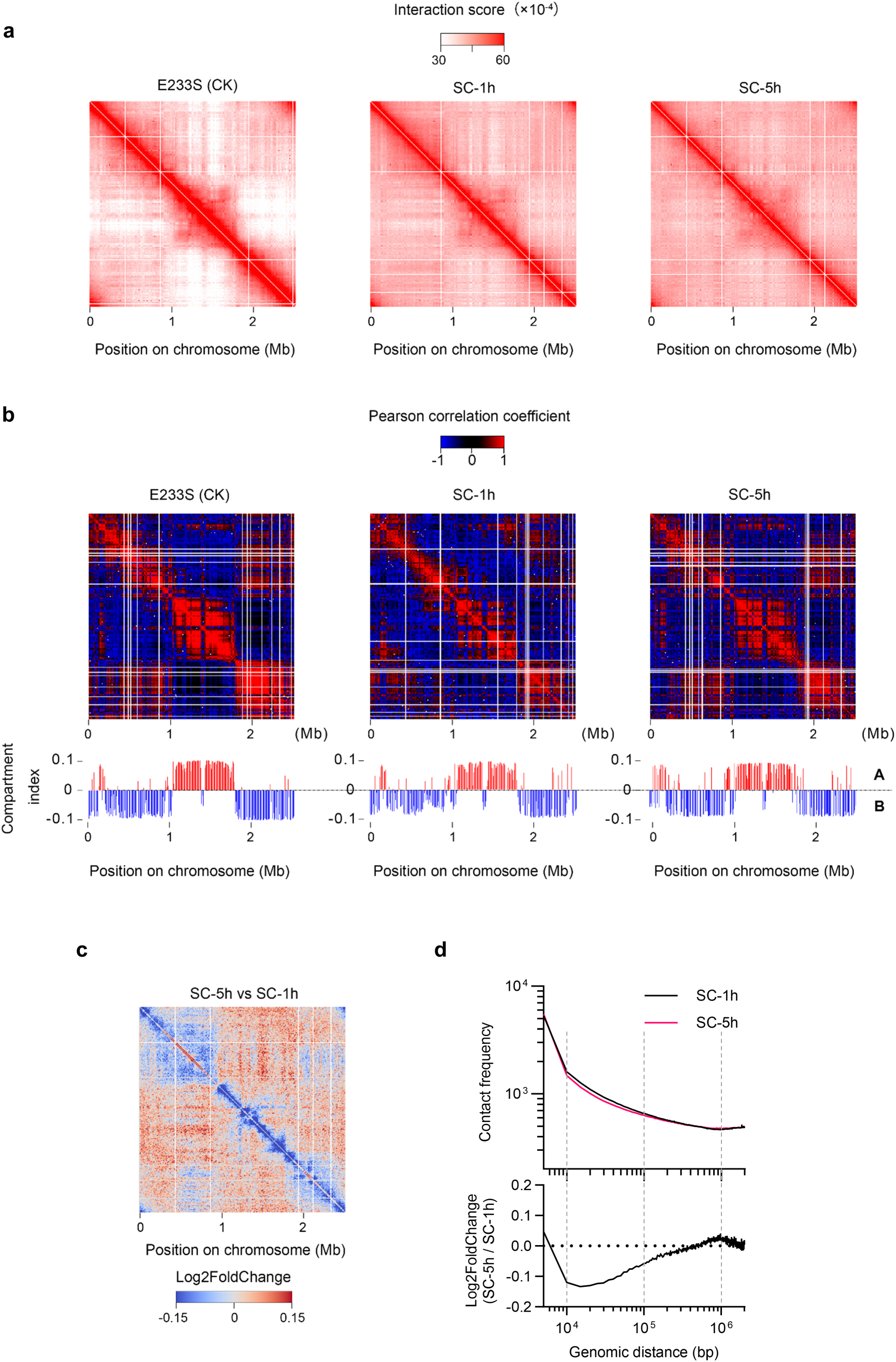
Analysis of chromatin compartmentalization in G2-synchronized cells during peak and trough transcription of *sul14a*. **a,** Hi-C contact matrices of synchronized cells 1 h (SC-1h) and 5 h (SC-5h) after release from G2 synchronization. Unsynchronized cells at exponential growth phase of E233S was used as the control (CK). The contact matrices represent interaction frequencies for pairs of 10-kb bins. Two biological replicates were combined for the analysis. **b**, Pearson correlation matrices of Hi-C data for E233S (CK), SC-1h, and SC-5h at 10-kb bin. **c**, Matrices of the ratios of Hi-C contacts between the SC-5h and the SC-1h samples. The data were analyzed at 10-kb resolution. The compartment index plots are shown at the lower panel. The compartment index plots are shown at the lower panel. **d,** Long-range chromosomal contacts increased in synchronized cells from 1 h to 5 h. Upper panels, the contact frequencies in SC-1h (black) and SC-5h (red) cells are plotted as a function of genomic distances (log scale; biological replicates combined). Bottom panels, Log2 fold changes in contact frequencies between comparative groups are shown against log-scaled genomic distances.

To further examine these changes, we generated a Hi-C contact matrix showing the contact ratio between SC-5h and SC-1h cells (Fig. 8c), and compared it with the corresponding matrix between pSeSD-Sul14a and pSeSD (Fig. 7c). Intriguingly, the interaction patterns during the trough-peak transition resemble those observed upon Sul14a overexpression, albeit to a less extent. Analysis of chromatin contact frequency distributions between trough and peak transcription of *sul14a* revealed that close- to mid-range interactions were attenuated, whereas long-range interactions increased (Fig. 8d). This trend parallels the pattern observed in Sul14a-overexpressing cells (Fig. 7d), though the magnitude of change was smaller under physiological conditions. It is noteworthy that long-distance interactions exhibit a more pronounced increase in Sul14a overexpression cells, a phenomenon potentially linked to compartment-scale architectural changes. These results imply that chromatin compartmentalization undergoes progressive modulation during the cell cycle, potentially coupled to the cyclic transcriptional changes of *sul14a*.

### Elevated Sul14a expression affects global transcription and its chromosomal localization

As Sul14a is globally enriched across chromatin (Fig. 6a,c), and its dynamic transcription during the cell cycle as well as its induced overexpression influence chromatin compartmentalization (Fig. 7a-c, Fig. 8a-c), we next analyzed how these changes affect the transcriptional landscape and their association with chromatin compartmentalization and Sul14a occupancy. We first compared the total transcript abundance between A and B compartments in the control and Sul14a-overexpressing cells. Consistent with previous report, the control cells exhibited significantly higher transcript abundance in A compartment than in B compartment (Fig. S7a and Fig. 9a) ^2^. While there was still a significant difference in abundance between A and B compartments in Sul14a-overexpressing cells, the magnitude of the difference was markedly reduced (Mann–Whitney U test, p = 0.0011). The results suggest that the transcription of genes in A compartment is reduced while that in B compartment is activated in general upon Sul14a overexpression. A similar trend, albeit weaker, was observed in the G2 synchronized cells during the trough and peak phases of *sul14a* transcription (Fig. S7b and 9b).

**Fig. 9.**
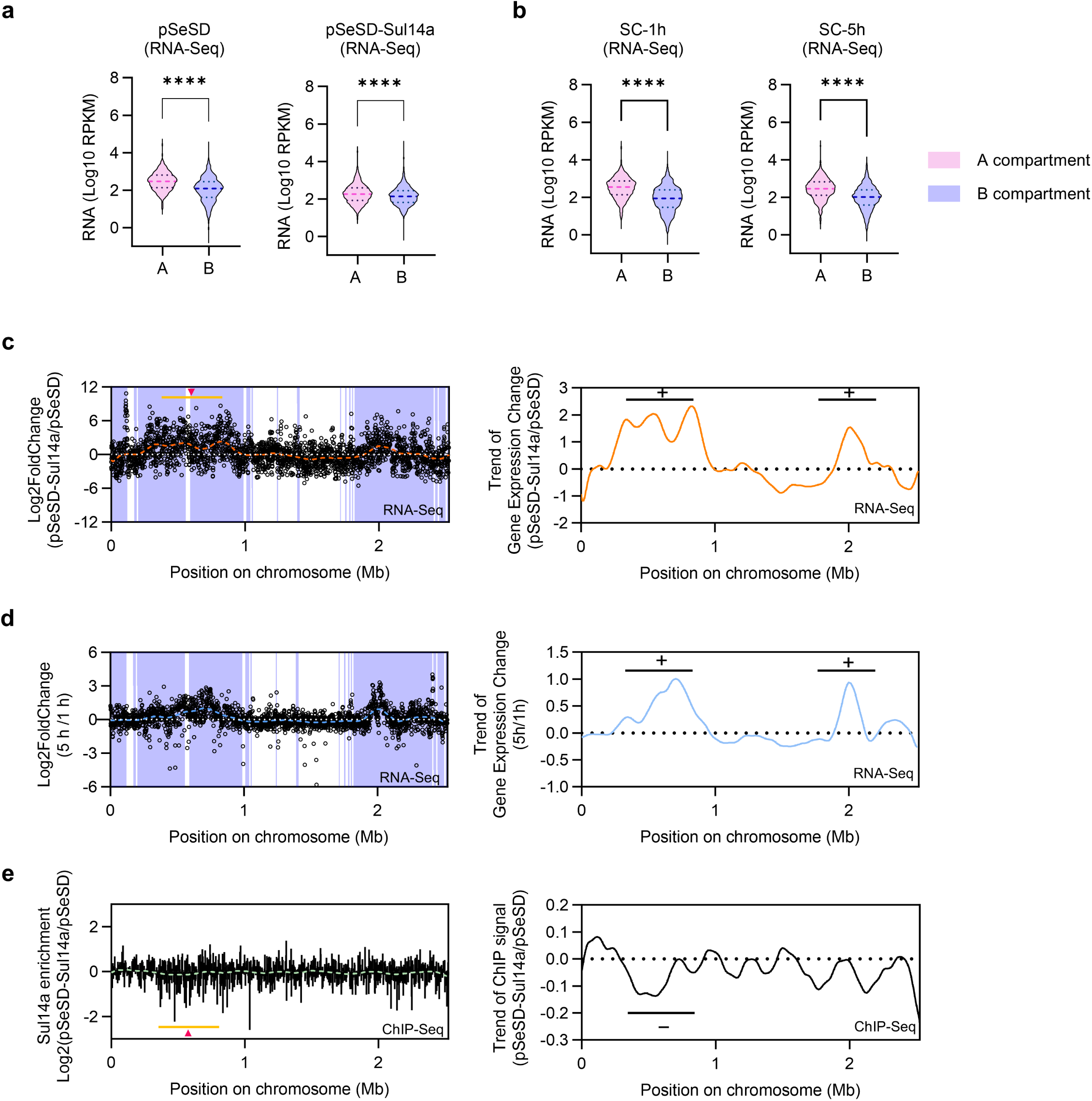
Elevated Sul14a expression affect global transcription and its chromosomal distribution. **a,** Violin plots showing the RNA transcript abundance (Log10 RPKM) in A/B compartments of the control and the Sul14a overexpression strain. Significance of the differences were tested using the two-tailed Student’s t-test with a threshold of *p* < 0.05 (\*\*\*\**P* < 0.0001). **b,** Violin plots showing the RNA transcript abundance (Log10 RPKM) in A/B compartments of 1h and 5h cells after release from G2 synchronization. **c**, Global transcriptional changes of protein-coding genes between cells carrying pSeSD and pSeSD-Sul14a. The log2 FPKM ratios (Log2Fold Change) were plot across the genome. The data were based on three biological replicates. Shaded lavender regions indicate the B compartment. The genome-wide trend of gene expression changes (right panel) were derived from LOWESS fitting employing a smoothing window of 20 data points. Genome coordinate is shown on the x-axis. **d**, Global transcriptional changes of protein-coding genes between cells at 5 h and 1 h after release from G2 synchronization ^8^. **e,** Genome-wide plot of the Log2FoldChange of the ChIP-seq signal (pSeSD-Sul14a/pSeSD) at a resolution of 2 kb. Genome coordinate is shown on the x-axis. The right panel shows the genome-wide trend of Log2FoldChange in the ChIP signal. Plus (+) and minus (−) indicate positions with concordant and discordant trends, respectively. The red triangles (panels c, e) indicate areas where there is a significant increase in transcriptional activity and decrease in occupancy of Sul14a.

We next analyzed the global gene transcriptional changes between Sul14a-overexpressing and control cells based on the transcriptomic data. The analysis revealed there is an extensive global reprogramming upon Sul14a overexpression, with more than 60% of annotated genes exhibiting > 2-fold transcriptional changes (padj < 0.05), 880 up-regulated and 794 down-regulated (Fig. S8a,b and Supplementary Data 2). When gene expression was analyzed in the context of chromatin compartments, Sul14a overexpression was associated with a general attenuation of transcription of A-compartment genes (36% down-regulated) and a marked increase in transcription of B-compartment genes (38% up-regulated) (Fig. 9c and S8c). These findings indicate a global shift in transcriptional activity consistent with the compartmental changes observed in Hi-C analyses.

To further evaluate whether this transcriptional pattern reflects cell-cycle associated *sul14a* dynamics, we compared transcriptomes of G2-synchronized cells sampled at the trough (1 h) and peak (5 h) of *sul14a* transcription using our published data ^8^. About 16% of genes in B compartment were up regulated, whereas the genes in A compartment showed only minimal transcriptional changes (Fig. 9d and Fig. S8d). Interestingly, the transcriptional trend between the peak and trough phases of *sul14a* expression mirrored those observed between Sul14a-overexpressing and control cells (Fig. 9c,d). Collectively, these findings suggest that elevated Sul14a levels preferentially enhance transcription of B-compartment genes while repressing A-compartment genes, linking Sul14a abundance to compartment-specific regulation of genome activity.

To further explore how Sul14a distribution responds to changes at expression level, we compared genome-wide ChIP-seq enrichment profiles between control and Sul14a-overexpressing cells. As shown in Fig. 9e, the difference profile exhibited multiple peaks and troughs across the chromosome, with a pronounced trough at approximately 0.5 Mb. Intriguingly, this reduction in Sul14a enrichment coincided with a region of elevated transcriptional activity within B compartment (0.2-1.0 Mb) (Fig. 9c). This inverse relationship suggests that increased Sul14a levels may induce local chromatin reorganization, thereby modulating its own binding distribution. It should be noted that the overall Sul14a enrichment appeared slightly reduced in the overexpression sample, which may partly reflect technical or biological factors such as altered chromatin accessibility or differences in crosslinking efficiency rather than a true decrease in binding. Taken together, these results reinforce the view that Sul14a functions as a chromatin organizer in *Sa. islandicus*, dynamically coordinating chromatin architecture and gene transcription.

## Discussion

Microbes of the order Sulfolobales have long served as valuable models for investigating the origins and molecular mechanisms underlying eukaryotic cellular features. One of the fundamental questions is how chromosomal structures and transcriptional dynamics are established and maintained. In this study, by integrating genetic, biochemical, ChIP-seq, transcriptomic, and Hi-C analyses, we demonstrate that a member of the Lrs14 family proteins functions as a chromatin-associated factor, whose cyclic expression affects chromatin compartmentalization and global transcription. Following the established nomenclature for NAPs in Sulfolobales, we designate the Lrs14 family protein, as Sul14a. Sul14a is well conserved within the Sulfolobales order and has a theoretical molecular mass of 14.3 kDa (Fig. S1). Several NAPs have been identified in Sulfolobales. Cren7 is conserved also in other orders of Crenarchaeota (now Thermoproteota) phylum, while Sul7d, Sul10a, and Sul12a are conserved in Sulfolobales ^12, 16, 28, 40^. Comparative analyses reveal that Sul14a shares key characteristics with known archaeal NAPs including high cellular abundance, non-specific DNA binding, dimer and oligomer formation ability, DNA bridging activity, and genome-wide chromatin enrichment, which supports its role as a NAP ^41^. Our transcriptomic data showed that all these NAPs genes, including *sul14a* and *cren7*, are transcribed in cyclic patterns in *Sa. islandicus* REY15A ^8^. Moreover, the *sul14a* gene appears to be essential, as we were unable to get a knockout mutant. Collectively, these findings strongly suggest that Sul14a plays a vital role in the establishment and maintenance of the dynamic chromatin structure in Sulfolobales.

As deletion of *sul14a* was unsuccessful, we constructed the Sul14a overexpression strains for phenotypic analysis. Overexpression of Sul14a exerted a pronounced impact on cell growth, chromatin compartmentalization, and cell cycle progress (Fig. 4b,c and Fig. 7a-c). Although induction (6h) of the protein may cause indirect effects, its overexpression exerted a strong and dominant impact, leading to cell cycle arrest at the G2 phase, so that potential indirect effects were not apparent. In addition, the overexpression of the DNA-binding-deficient mutants did not reproduce the effects induced by wild-type protein overexpression, indicating that the DNA binding activity of Sul14a is crucial for its cellular function (Fig. 4b,c). Notably, while *sul14a* transcript levels increased by approximately 25- to 40-fold in the overexpression strain, the corresponding protein abundance increased only about threefold (Fig. S5a-d), implying that Sul14a is subjected to additional layers of regulation, possibly at the post-transcriptional or post-translational level.

It is intriguing that although *sul14a* transcription exhibits a clear cyclic pattern, the Sul14a protein level remains relatively stable throughout the cell cycle (Fig. 1d). Comparison of chromatin compartmentalization and global transcription between Sul14a-overexpressing cells and synchronized cells at the peak and trough phases of *sul14a* transcription revealed similar overall trends (Figs. 7-9). These observations suggest that during the peak transcription of *sul14a*, a larger fraction of Sul14a is bound to chromatin, whereas its chromatin association is reduced during the trough phase. We propose that post-translational modifications (PTMs), such as phosphorylation, may regulate the DNA-binding activity of Sul14a. Previous studies have shown that phosphorylation modulates the enzymatic activity of the Holliday junction resolvase Hjc and the promoter-binding activity of the DNA damage regulator Orc1-2 ^42, 43^. A similar mechanism may apply to Sul14a. As the *Sa. islandicus* kinase ePK2 is expressed in a cell cycle-dependent manner ^7^, it could potentially target conserved serine and threonine residues located near the DNA-binding HTH domain of Sul14a (Fig. 1a). Future studies should determine whether phosphorylation or other PTMs, including methylation and acetylation ^44, 45^ modulate Sul14a’s activity. In addition, recent work from our group has identified that the *sul14a* promoter is a target of one of the aCcr transcriptional regulators aCcr3 (archaeal cell cycle regulator 3) ^11^, suggesting that both transcriptional and post-translational mechanisms may contribute to the multilayered regulation of the function of Sul14a.

Two previous studies have characterized Sul14a on its *in vitro* biochemical properties ^23, 24^. The Sul14a protein in *Sa. solfataricus* is identical in amino acid sequence to that of *Sa. islandicus* REY15A, the strain used in this study, reenforcing close phylogenetic relationship between these two species and the conservation of this protein. In early work, this protein was termed Sta1 (Sulfolobus transcription activator 1), due to its role in activating transcription of a viral gene ^23^ and the *radA* paralog gene ^24^. These findings indicate that Sta1/Sul14a possesses non-specific DNA-binding activity with an AT-rich preference and functions as a transcriptional activator in certain contexts. Consistent with these observations, our results demonstrate that Sul14a binds DNA non-specifically, bridges DNA molecules *in vitro*, and is involved in global transcription through chromatin organization. Together, these findings suggest that Sul14a participates in both promoter-based and chromatin organization-based transcriptional regulation activities.

As reported, all six *lrs14*-like genes in *S. acidocaldarius* could be deleted or disrupted, with three clean deletions (*saci_0102, saci_0446, saci_1242*) and three insertional disruptions (*saci_0133*, *saci_1219*, and *saci_1223*). All mutants remained viable with normal growth ^46^. The failure to obtain a viable *sul14a* (homolog of *saci_0102*) deletion mutant in *Sa. islandicus* may be attributed to species-specific differences in genetic background or in the essentiality of nucleoid-associated proteins. While *S. acidocaldarius* and *Sa. islandicus* are closely related members of the order Sulfolobales, they display notable differences, for example, in chromosomal organization ^2^ and cell division mechanism ^5, 47^, which may contribute to the differential essentiality of certain genes between species. Therefore, it is possible that Saci_0102 performs a nonessential yet supportive role in *S. acidocaldarius*, but Sul14a is indispensable for chromatin integrity or cell cycle progression in *Sa. islandicus*.

Based on our findings, we propose a model for Sul14a functioning *in vivo* (Fig. 10). As an essential and highly abundant nucleoid-associated protein, Sul14a binds and bridges DNA to maintain genomic stability and higher-order chromosome architecture. Through its association with chromatin, Sul14a exerts global effects on gene transcription by modulating chromatin accessibility and compartmentalization. At local levels, Sul14a binds preferentially to TSS-proximal regions, thereby influencing promoter activity and then regulates gene expression. Its transcription level increases from G2 phase to late S phase, regulating global gene expression and modulating chromatin structure during the cell cycle. These regulatory activities collectively result in repression of genes in A compartment and activation of those in B compartment. However, excessive Sul14a disrupts chromatin conformation and compartmental integrity, leading to G2-phase arrest. Our work identifies Sul14a as a global chromatin regulator that combines both local transcriptional modulation and higher-order chromosomal organization activities.

**Fig. 10.**
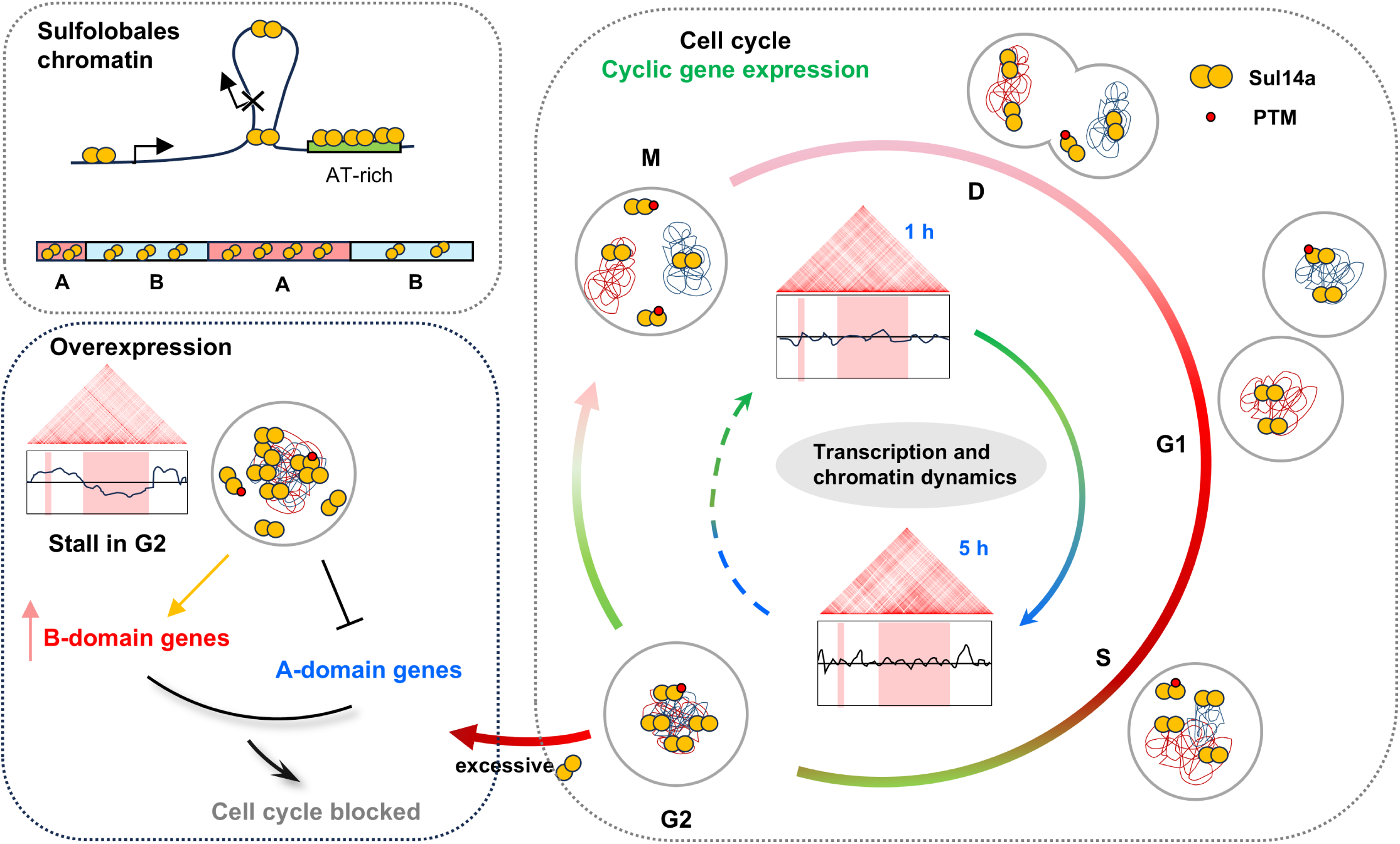
A model illustrating the role of Sul14a in cell cycle regulation and chromosome organization. Sul14a (orange balls) binds and bridges DNA to maintain genomic stability and overall chromosome conformation. It preferentially binds AT-rich DNA, influencing promoter activity and regulates gene expression. Cell cycle-dependent upregulation of *sul14a* (G2→late S phase, green to dark red) results in dynamic remodeling of the 3D genome architecture and triggers changes of the transcriptome, orchestrating cell cycle phase-specific gene expression. Excessive Sul14a silences A compartment genes and activates the B compartment genes, leading to chromatin disorganization and cell cycle arrest in the G2 phase. PTM (red balls), post-translational modification which is postulated to be involved in the regulation of the DNA binding activity of Sul14a.

In summary, this study establishes that Sul14a is a novel cell cycle-related NAP in Sulfolobales. Future studies should aim to elucidate the molecular mechanisms underlying Sul14a-mediated chromatin modulation and to explore the functional interactions between Sul14a and other chromatin proteins including Cren7, Sul7d, Sul10a, and Sul12a. Given the presence of multiple Lrs14 family proteins in *Sa. islandicus* (Table S1 and Fig. S9) and related species, it is plausible that other members also act as NAPs with complementary or specialized functions. Investigating the individual and cooperative functions of these proteins, both independently and in conjunction with Sul14a, will provide valuable insights into their contributions to cell cycle regulation and responses to environmental cues.

## Methods

### Strains and growth conditions

All *Sa. islandicus* strains were derived from *Saccharolobus islandicus* REY15A wild type strain. Cultivation was performed in STV medium (basal salt supplemented with 0.2 % [w/v] sucrose (S), 0.2 % [w/v] tryptone (T), and a mixed vitamin solution (V)) or ATV medium (basal salt with 0.2 % D-arabinose (A) instead of sucrose), adjusted to pH 3.0 using sulfuric acid. Uracil (50 μg/mL) was added to complement the biosynthetic auxotrophy of the *ΔpyrEFΔlacS* parent strain E233S (hereafter E233S). Solid medium was prepared using gelrite (0.8 % [w/v]) by mixing 2 × STV with an equal volume of 1.6 % gelrite. Other ingredients and media preparation are according to Yang et. al. ^8^. All plates were incubated at 75°C for 5–7 days for single colonies and liquid cultures were cultivated aerobically at 75°C with 150 rpm orbital agitation.

### Purification of recombinant Sul14a

*Sa. islandicus* Sul14a proteins were expressed with a C-terminal 6 × His-tag in *E. coli* BL21 (DE3) cells. The *sul14a* gene and its mutant variants were amplified by PCR and cloned into pET22b plasmid digested by *NdeI* and *SalI*. Recombinant plasmids were transformed into *E. coli* by heat shock. Cultures were grown in LB at 37°C to an OD_600_ = 0.4-0.6 and induced with 1 mM IPTG at 37°C for 4 h. The cells were collected by centrifugation at 7,000 × *g* for 10 min and resuspended in buffer A (50 mM Tris-HCl, pH 6.8, 50 mM NaCl, and 5% glycerol). After sonication, the supernatant was obtained by centrifugation at 12000 *g* for 20 min and then heat-treated at 70°C for 30 min to denature *E. coli* proteins. The heat-stable fraction was first purified over Ni-NTA agarose (Invitrogen) pre-equilibrated with buffer A and bound proteins were eluted with buffer A containing 400 mM imidazole. Further purification and estimation of the protein oligomeric state were performed using a size exclusion chromatography (SEC) column (Superdex 200 Increase 10/300 GL, GE Healthcare). The protein concentration was determined using the Bradford method.

### Generation of Sul14a overexpression strains

The overexpression strains were generated based on the shuttle vector pSeSD as previously described ^48^. The *sul14a* gene and its mutant derivatives were amplified by PCR with a C-terminal 6 × His-tag and then cloned into pSeSD plasmids digested by *NdeI* and *SalI* and transformed into E233S by electroporation. The non-tagged *sul14a* was also retained. Sequencing was used to verify the presence of the correct point mutation(s). Strains containing pSeSD (empty vector), pSeSD-Sul14a and pSeSD-Sul14a-C-His (for the wild type overexpression), pSeSD-Sul14a-R80A/K82A/K87A/R90A-C-His (for the overexpression of DNA binding deficient mutants) were generated. The strains and plasmids constructed and used in this study are listed in Table S2 and the oligonucleotide sequences for PCR are listed in Table S3. The growth curves of Sul14a overexpression strains were obtained from an initial OD_600_ of 0.03-0.05, and samples were taken every 6 or 12 h to get the OD_600_ in STV or ATV medium. For RNA-Seq, Hi-C and ChIP-seq samples, strains containing pSeSD and pSeSD-Sul14a were grown to approximately OD_600_ = 0.2 in STV medium. D-arabinose was then added to cultures to a final concentration of 0.2% for induction. All induced samples were collected 6 h post-induction. Overexpression of Sul14a was confirmed by Western blotting.

### Cell cycle synchronization

*Sa. islandicus* REY15A cells were synchronized as previously described with slight modifications. The parent strain E233S and cells carrying pSeSD (empty vector) or pSeSD-Sul14a were cultivated aerobically at 75°C with 150 rpm orbital agitation in 20 mL STV (U) medium. When cultures reached OD_600_ 0.5–0.7, cells were transferred into 50 mL fresh medium with an initial estimated OD_600_ of 0.05. At OD_600_ 0.18–0.2, 6 mM acetic acid (Sigma-Aldrich) was added to induce G2-phase arrest over 6 h. For cells containing pSeSD-Sul14a, 0.2% D-arabinose was added 2 h before release. Cells were then centrifuged at 3,000 × *g* for 10 min at room temperature and washed twice with 0.7% (w/v) sucrose and resuspended in 50 mL of pre-warmed ATV medium and cultivated as above for subsequent analysis. For E233S, cells were cultured in STVU medium all the time.

### Gene expression analysis by quantitative reverse transcription PCR (qRT-PCR)

The total RNA of the indicated cells was extracted by SparkZol Reagent (SparkJade, Shandong, China) following the manufacturer’s instructions. First-strand cDNA was synthesized by reverse transcription of extracted RNA using Evo M-MLV RT Mix Kit (Accurate Biotechnology, Hunan, China). The qRT-PCR was performed on the CFX96 Touch™ Real-Time PCR System (Bio-Rad, Hercules, CA, USA) using the SYBR Green Premix Pro Taq HS qPCR Kit (Accurate Biotechnology, Hunan, China) to evaluate the mRNA levels of target genes. The two-step PCR amplification standard procedure was as follows: 95 °C for 30 s, followed by 40 PCR cycles consisting of 95 °C for 5 s and 60 °C for 30 s. The relative expression level of the target gene was calculated using the comparative threshold cycle (CT) method (2^−ΔΔCT^) and the 16S rDNA was used as an endogenous control gene. The primers for qRT-PCR are listed in Table S3.

### Flow cytometry

The procedure for the flow cytometry of *Sa. islandicus* was conducted as previously described ^8^. *Sa. islandicus* cultures (0.3 mL aliquots) were fixed in 70 % ice-cold ethanol and stored at 4 °C. The fixed cells were pelleted at 800 × *g* for 20 min, washed twice with 1 mL PBS buffer and resuspended in 150 μL PBS buffer. Nucleic acids were stained with SUPER Green I (Fanbo Biochemicals, Beijing, China) or 50 μg/mL propidium iodide (PI) (Sigma Aldrich). The stained samples were analyzed using the ImageStreamX MarkII Quantitative imaging analysis flow cytometry (Merck, Darmstadt, Germany) equipped with a 488 nm laser. A dataset of at least 200,000 cells was obtained for each cell sample and analyzed with FlowJo.

### Western blotting

The antibody against TBP was prepared utilizing synthetic specific peptides (amino acids 18-31, SIPNIEYDPDQFPG for TBP) and the antibody against Sul14a was prepared utilizing pure protein expressed and purified from *E. coli*. The antibodies were produced in rabbits by HuaAn Biotechnology, Hangzhou, China (for TBP) or Mabnus, Wuhan, China (for Sul14a). *Sa. islandicus* cells were pelleted at 7,000 × *g* for 10 min and resuspended in buffer A (50 mM Tris-HCl pH 8.0, 300 mM NaCl and 5% glycerol). The samples were supplemented with 5× loading buffer and boiled for 10 min. Proteins were fractionated on 15 % SDS-PAGE and transferred to polyvinylidene difluoride (PVDF) membranes (200 mA, 2 h, 4 °C). Membranes were blocked with 5 % skim milk (2 h, RT), incubated with primary anti-rabbit antibody (Anti-Sul14a) and the secondary goat anti-rabbit HRP conjugate antibody (TransGen Biotech, Beijing, China) according to the manufacturer’s instructions. The hybridization signals were visualized on Amersham ImageQuant 800 (Cytiva, USA). The protein relative levels were analyzed and quantified based on the grayscale values of Western blotting figures by ImageJ. Sul14a band intensities were normalized against an internal standard TBP. Besides, the estimated copy number of Sul14a per cell was calculated from a linear regression of a standard curve generated from purified Sul14a. The mean and SEM of at least two replicates for each condition are reported.

### Electrophoretic mobility shift assay (EMSA)

Substrates used in EMSA experiments were generated by annealing the complementary oligonucleotides with 5′ FAM-labeled oligonucleotides or by PCR with 5’-FAM-labeled primers (Table S3). The reaction with 10 nM 200bp-FAM-labeled dsDNA as substrate was taken at different concentrations of Sul14a and its mutants (0, 0.05, 0.1, 0.2, 0.4, 0.8, 1.6, and 3.2 µM), and the reaction system was 50 mM Tris-HCl pH 6.8, 25 mM NaCl, 10 mM MgCl_2_, 5% glycerol. The reaction with 20nM 55bp-FAM-labeled AT/GC substrates was added with another Sul14a concentration gradient (0, 0.1, 0.2, 0.4, 0.8, 1.6, 3.2, 6.4, and 9.6 µM). After 30 min incubation at room temperature, complexes were resolved on 6% or 8% polyacrylamide gels with 0.5 × TBE buffer at 120 V for 60 min. Fluorescent signals were captured using an Amersham ImageQuant 800 (Cytiva, USA). All DNA binding experiments were performed at least in triplicate.

For the reaction with plasmid DNA as substrate, different concentrations of Sul14a (0, 0.05, 0.1, 0.2, 0.4, and 0.8 μM) were taken with 300 ng of plasmid pBR322 with different types (supercoiled, linearized, and nicked), and the reaction system was 50 mM Tris-HCl pH 6.8, 25 mM NaCl. The reaction was incubated at room temperature or 37°C for 30 min. After incubation, the results were detected by 1% agarose gel electrophoresis with 0.5×TAE buffer at 110 V, 40 min, and visualized by EC3 Imaging System (Ultra-Violet Products Ltd, Cambridge UK).

### DNA bridging assay

The DNA bridging assay was performed as previously described with minor modifications ^36, 37^. In short, the biotinylated 1,000 bp bait DNA was prepared by PCR using a 5′ biotin-labeled forward primer and a reverse primer (Table S3), and 500 bp prey DNA was generated by PCR with a 5’-FAM-labeled primer. For each bridging assay measurement, the procedure was as follows: 20 μL of streptavidin magnetic beads (BEAVERBEADS, China) were washed three times in binding buffer (20 mM Tris-HCl pH 7.5, 50 mM KCl, 5 mM EDTA, 1 mM DTT, 5% glycerol, and 40 μg/mL bovine serum albumin (BSA)), incubated with 200 ng bait DNA for 15 min at room temperature. The beads were collected by magnet and washed three times. A concentration of 15 nM of prey DNA and the indicated amount of Sul14a (0, 2.5, 5, 10, and 15 μM), and binding buffer were added in a total volume of 40 μL and incubated for 30 min at 20 °C. After collection, the beads were washed twice with binding buffer and once with BSA-free binding buffer, and the beads were then resuspended in 30 μL BSA-free binding buffer. Sul14a was subsequently removed from the reaction by adding 10 μg/μL proteinase K and 0.5% SDS and treating the beads for 30 min at 65°C. Beads were then collected by magnet and the supernatant was collected and resolved on a 6% or 8% native polyacrylamide gel (0.5 x TBE, 120 V, 60 min), the fluorescent signals were captured using an Amersham ImageQuant 800 (Cytiva, USA).

For plasmid DNA, the nicked substrates were generated by digesting pBR322 with Nb. *Bpu10I* (Thermo Fisher Scientific), while linear forms were obtained by digesting pBR322 with *BamHI*. Nicked, supercoiled, linearized pBR322 (500 ng) was incubated with indicated amount of Sul14a (0, 5, 10, 15, and 20 μM), processed as above, the final supernatant was resolved on a 0.7% agarose gel (110 V, 90 min), and visualized by EC3 Imaging System (Ultra-Violet Products Ltd, Cambridge UK). All DNA bridging experiments were performed at least in triplicate.

### ChIP-Seq and data analysis

Chromatin immunoprecipitation (ChIP) was performed as described ^8, 49^. Cells carrying pSeSD or pSeSD-Sul14a were collected following the workflow outlined in Fig. 3a. Cultures (300 mL) were cooled to room temperature, crosslinked with 1% formaldehyde (37% stock) and quenched with 125 mM glycine. Samples were pelleted (7,000 × *g*, 10 min), washed with PBS, and resuspended in 1.2 mL TBS-TT buffer (20 mM Tris-HCl pH 7.5, 150 mM NaCl, 0.1% Tween-20, 0.1% TritonX-100). The resuspensions were sonicated on ice with a JY92-II sonicator (Scientz Biotechnology, Ningbo, China) to yield 200-1,000 bp DNA fragments. After centrifugation (6,000 × g, 10 min, 4 °C), 100 µL supernatant was used as input. Immunoprecipitation samples were incubated with 50 µL purified anti-Sul14a antibody for 2 h (4°C, rotation), followed by the addition of 50 µL protein A Sepharose beads (Cytiva) and continued rotation for 4 h at 4°C. The immune complexes were captured by the protein A-Sepharose and washed 5 times with TBS-TT buffer. Then the beads were washed once with TBS-TT containing 500 mM NaCl and once with TBS-TT containing 0.5 % Tween 20 and Triton X-100. The DNA-Sul14a complexes were eluted by incubating the beads with the elution buffer (20 mM Tris-HCl pH 7.5, 10 mM EDTA, 0.5 % SDS) in a total volume of 200 µL at 65 °C for 30 min. Sul14a was removed through proteinase K digestion (10 µg/mL) at 65 °C for 6 h, followed by 37 °C for 10 h. The immunoprecipitated DNA was then purified using a DNA Cycle-Pure Kit (Omega Bio-tek, Norcross, USA). ChIP reactions were performed in replicates and pooled. ChIP DNA degradation and contamination were monitored on agarose gels. DNA purity was checked using the NanoPhotometer spectrophotometer (IMPLEN, CA, USA). The purified DNA was used for ChIP-seq library preparation (constructed by Personalbio Corporation, Nanjing, China). Library quality was assessed on the Agilent Bioanalyzer 2100 system. Subsequently, pair-end sequencing of the sample was performed on the Illumina platform (Illumina, CA, USA).

Reads from single or multiple biological replicates were mapped to the genome of *Sa. islandicus* REY15A (GeneBank ID: CP002425.1) using Bowtie 2 v2.5.4 with default parameters ^50^. To calculate protein enrichment on the chromosome, ChIP-seq coverage was divided by input coverage of the region after normalizing the total number of reads mapped to the chromosome. After mapping, ChIP reads and input reads were compared using bamCompare (parameters: -bs 10, –scaleFactorsMethod readCount). The peak calling was done utilizing the MACS3 to identify regions of IP enrichment over the background. Metagene analyses on RefSeq protein-coding genes were performed using functions implemented in deepTools v3.5.5 ^51^.

### RNA-Seq and data analysis

The transcriptomic analysis was performed as previously described ^8, 52^. Strains carrying pSeSD and pSeSD-Sul14a were collected after 6 h induction according to the method (“Generation of Sul14a overexpression strains”). The pellet was washed in PBS, frozen with liquid nitrogen and transported on dry ice. Total RNA was extracted using the Trizol reagent (Ambion, Austin, TX, USA) and assessed by Agilent 2100 bioanalyzer. The samples were analyzed and sequenced by Novogene Bioinformatics Technology (Beijing, China).

Reads from three biological replicates were pooled and mapped to the genome of *Sa. islandicus* REY15A using Bowtie 2 v2.5.4 ^50^. After the removal of multi-mapping reads, the remaining reads were processed using bamCoverage implemented in deepTools v3.5.5 ^51^ to calculate Reads Per Kilobase region per Million mapped reads (RPKM) for genomic bins of fixed size. To quantify expression levels for protein-coding genes and noncoding RNA, reads from each biological replicate were mapped using Salmon v0.8.2 ^53^. When calculating the log2 fold change to determine differentially expressed genes, the data generated by Salmon was further processed using DEseq2 (|log2FC| > 1, FDR < 0.05).

### Hi-C

The method for Hi-C sample preparation was performed as previously described ^54, 55^ with slight modifications. For the cells carrying pSeSD and pSeSD-Sul14a plasmids, the samples were collected as Fig. 3a. For the synchronized E233S, cells were collected at 1 h and 5 h after release. Briefly, for each Hi-C experiment, 5 billion cells were cross-linked with 1% formaldehyde (30 min, 25°C), quenched with 0.25 M glycine (10 min, 25°C) and pelleted (6,000 × g, 10 min, 4°C). The pellets were washed twice with PBS and stored at −80°C until use.

The frozen pellets were resuspended and diluted to an OD_600_ of 4 in 1 × NEBuffer 2, and 400 μL of the cell suspension was centrifuged (21,000 × g, 4 °C,5 min). The pellets were then resuspended in 50 μL of 1 × NEBuffer 2 and lysed with 5.55 μL of 10% SDS at 65°C for 15 min with agitation. Lysates (37.5 μL) were cooled on ice (90 s), then mixed with 11.2 µL of 10× NEBuffer 2 and 30 µL of 20% Triton X-100. Chromosomal DNA was digested by adding 15 μL of 100 U/μL *HindIII* (NEB) and incubating for 4 h at 37°C. After heat inactivation of *HindIII* (65°C, 20 min), the *HindIII* DNA ends (50 μL) were filled and labeled with biotin by adding 10 μL of 0.4 mM biotin-14-dCTP (Thermo), 2 μL each of 2 mM dATP, dTTP, dGTP, 1.8 μL of 10 × NEBuffer 2, and 0.5 μL of 5 U/μL Klenow Large Fragment (NEB), with incubation at 20°C for 30 min. Additionally, for the sample of E233S cells, the chromosomal DNA was digested by *Sau3AI* (NEB). The digestion and fill-in reactions were quenched by adding 7.7 μL of 10% SDS and 1.4 μL of 0.5 M EDTA (pH 8.0) at 21°C for 5 min. After ligation at 16°C overnight using T4 DNA ligase (NEB), the samples were reversely cross-linked by adding 100 μL of 10% SDS, 50 μL of 0.5 M EDTA (pH 8.0), and 5 μL of 20 mg/mL proteinase K with incubation at 65°C for 6 h, followed by further incubation at 37°C for more than 6 h. The first DNA purification step was completed with a DNA Cycle-Pure Kit (Omega Bio-tek, Norcross, USA). The DNA was dissolved in 40 μL of 1 × NEBuffer 2 containing 0.1 mg/mL RNase A and incubated at 37°C for 30 min. Subsequently, 30 μL of DNA obtained from the above purification was mixed with 4.5 μL of 10 × NEBuffer 2, 0.375 μL of 20 mg/mL BSA (Thermo Fisher Scientific), 3.75 μL of 2 mM dCTP, 0.375 μL of 3U/μL T4 DNA Polymerase (NEB), and 36 μL of Milli-Q water to a total volume of 75 μL and then incubated at 20°C for 4 h to remove biotin from unligated free DNA ends. The second DNA purification was immediately followed by using a DNA Cycle-Pure Kit (Omega Bio-tek, Norcross, USA). DNA was sheared for fragmentation.

Sheared DNA (55.5 μL) was used to construct the Hi-C library with NEBNext Ultra DNA Library Prep Kit for Illumina and NEBNext Multiplex Oligos for Illumina. End repair, adaptor ligation, and size selection were performed according to the manufacturer’s instructions. The purified DNA was subjected to biotin purification with Dynabeads MyOne Streptavidin C1 (Thermo Fisher Scientific). Before use, 10 μL of beads per sample were washed 4 times with B&W Buffer (5 mM Tris-HCl pH 7.5, 0.5 mM EDTA, 1M NaCl) and resuspended in 135 μL of B&W Buffer. The bead suspension (135 μL) was added to each DNA sample and rotated at 21°C for 30 min. The beads were then washed 3 times with B&W Buffer and once with 1 × NEBuffer 2 (without DTT). Each sample was resuspended in 15 μL of 0.1 × TE (pH 8.0) and amplified by PCR in a 50 μL reaction for 14 cycles. The PCR products were purified with AMPure XP Beads (Beckman Coulter) and paired-end sequenced (150 bp x 2) on the DNBSEQ-T7 platform at ANOROAD GENOME Technology, Beijing, China. Each Hi-C experiment was performed with two biological replicates.

### Hi-C data analysis

The raw Hi-C reads were mapped and processed using HiC-Pro 3.1.0 ^56^. Reads were mapped to the genome of *Sa. islandicus* REY15A with default parameters. Valid read pairs were used to generate raw Hi-C contact matrices and the iced matrices were visualized using Treeview v1.2.1. The genome was binned at 5-kb for samples pSeSD and pSeSD-Sul14a and 10-kb for samples CK, SC-1h and SC-5h. When combining biological replicates, valid read pairs from them were pooled before generating raw Hi-C matrices. Biological replicates were analyzed separately to assess data reproducibility. Pearson correlation matrices and compartment index plots were generated as described previously using HiTC ^2, 57^. To compare contact maps between conditions, we first normalized the maps and then calculated the ratio for each bin by dividing the contact counts in one condition by those in the other. The resulting ratios were log2-transformed and visualized using a blue-to-red color scale. The comparison between multiple Hi-C matrices of the Hi-C counts enrichment at different genomic ranges/distances up to the whole chromosome was performed using HiCExplorer 3.7.3 ^58^.

### Data analysis

The gene compartment assignment was based on the genomic location of each gene’s midpoint. For the comparative analysis of ChIP-seq enrichment profiles and RNA transcript abundance between A/B chromatin compartments, statistical significance was evaluated using the two-tailed Student’s t-test with a significance threshold of *p* < 0.05. If needed, additional quantification and statistical analyses were performed using R software (http://www.R-project.org).

### Statistics and reproducibility

The specific number of replicates employed in each experiment is provided in the corresponding methods and figure legends. All quantitative data was expressed as the mean ± standard error (SEM) and analyzed using the GraphPad Prism 10.1 statistic software (GraphPad Software). The Kd and SEM values for DNA binding in Fig.2 and Fig.S3 were obtained through nonlinear regression analysis using a one-site specific binding model. For Fig. 6b and 6d, correlation analysis was conducted using Spearman’s correlation and Gaussian fitting was applied to perform the curve fits. For Fig.9a,b and Fig.S5d,e, the statistical analysis was performed using unpaired *t* test, statistically significant differences are denoted in graphs with * standing for *p*-value, where * refers to *p* < 0.05, ** to *p* < 0.01, *** to *p* < 0.001, and **** to *p* < 0.0001. For Fig. 9, LOWESS fitting was performed employing a smoothing window of 20 data points.

### Data availability

All sequencing data used in this study have been deposited in the NCBI Sequence Read Archive (SRA) under project number PRJNA1245118. The RNA-seq data for the synchronization samples were previously reported in ref. 8, and is available in the GEO, accession number GSE220819. Supplementary Data 1 provides the number and locations of peaks from the ChIP-seq data. Supplementary Data 2 provides the results of changes in gene expression levels from the transcriptome. All source data used to produce the graphs in this work are provided in Supplementary Data 3. Uncropped blots/gels are provided in Supplementary Data 4. The DNA sequence information in this study is provided in Supplementary information.

### Code availability

No custom code was generated for this work.

## Acknowledgements

The work was funded by the National Natural Science Foundation of China (Nos. 32393973 and 32370033 to Y. S., 31970119 to J.N., and 31900055 to Q. H.), the National Key R&D Program of China (Grant No. 2020YFA0906800 to J. N. and Q. H.), and the State Key Laboratory of Microbial Technology Open Projects Fund (M2023-20, M2025-12) to Y.S.. We thank all the lab members of the CRISPR and Archaea Biology Research Centre for helpful discussions and technicians from the Core Facilities for Life and Environmental Sciences, State Key Laboratory of Microbial Technology of Shandong University for assistance. We thank Guannan Lin, Zhifeng Li, Jing Zhu, and Jingyao Qu of the Core Facilities for Life and Environmental Sciences, State Key laboratory of Microbial Technology of Shandong University for flow cytometry analysis (ImageStreamX MarkII).

## Contributions

Conceptualization: Yulong Shen, Qi Gan, and Haodun Li. Methodology: Qi Gan, Haodun Li, Qihong Huang, and Yunfeng Yang. Investigation, data analysis and visualization: Qi Gan and Haodun Li. Writing—original draft: Qi Gan and Yulong Shen. Funding acquisition: Yulong Shen, Jinfeng Ni and Qihong Huang. Review and editing: all authors.

