## Supplementary for "An Lrs14 family protein functions as a nucleoid-associated protein regulating cell cycle progression in Sulfolobales"

This file includes:

Supplementary Figures: Fig. S1 to S9

Supplementary Tables: Table S1 to S3

Supplementary Materials and Methods

The DNA sequence information in the paper

Legends for the supplementary data

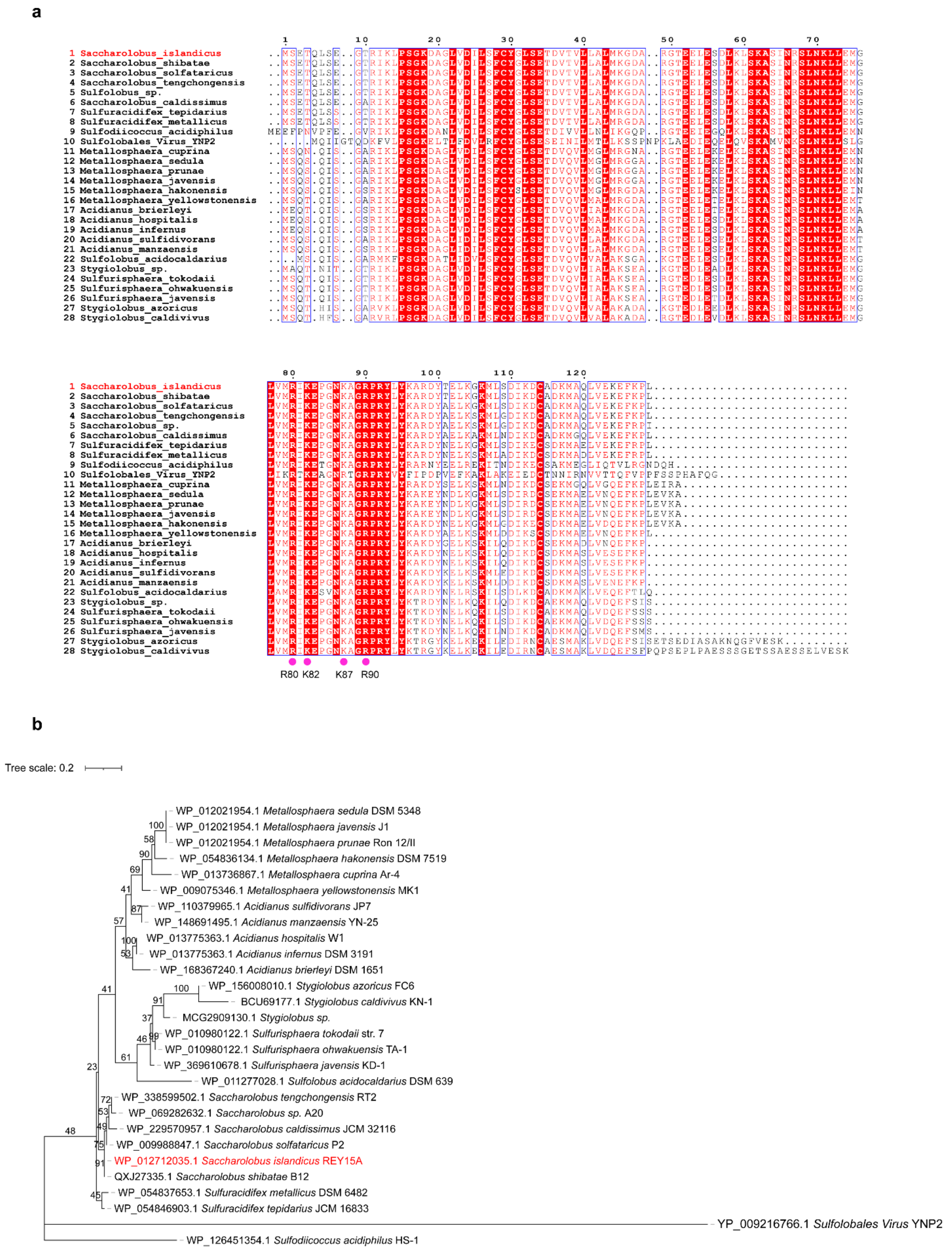


**Fig. S1. Sul14a is conserved in Sulfolobales.**

**a,** Multiple sequence alignment of Sul14a homologs from 28 *Sulfolobus* species. The alignment was generated using Clustal Omega, with conserved DNA-binding residues highlighted at the bottom. **b**, Phylogenetic tree of Sul14a homologs based on the sequence alignment in (a). The tree was generated by Maximum Likelihood method with the LG+G model by IQ-Tree 2. The scale bar refers to the phylogenetic distance. Bootstrap values (1000 replicates) are shown next to branches.

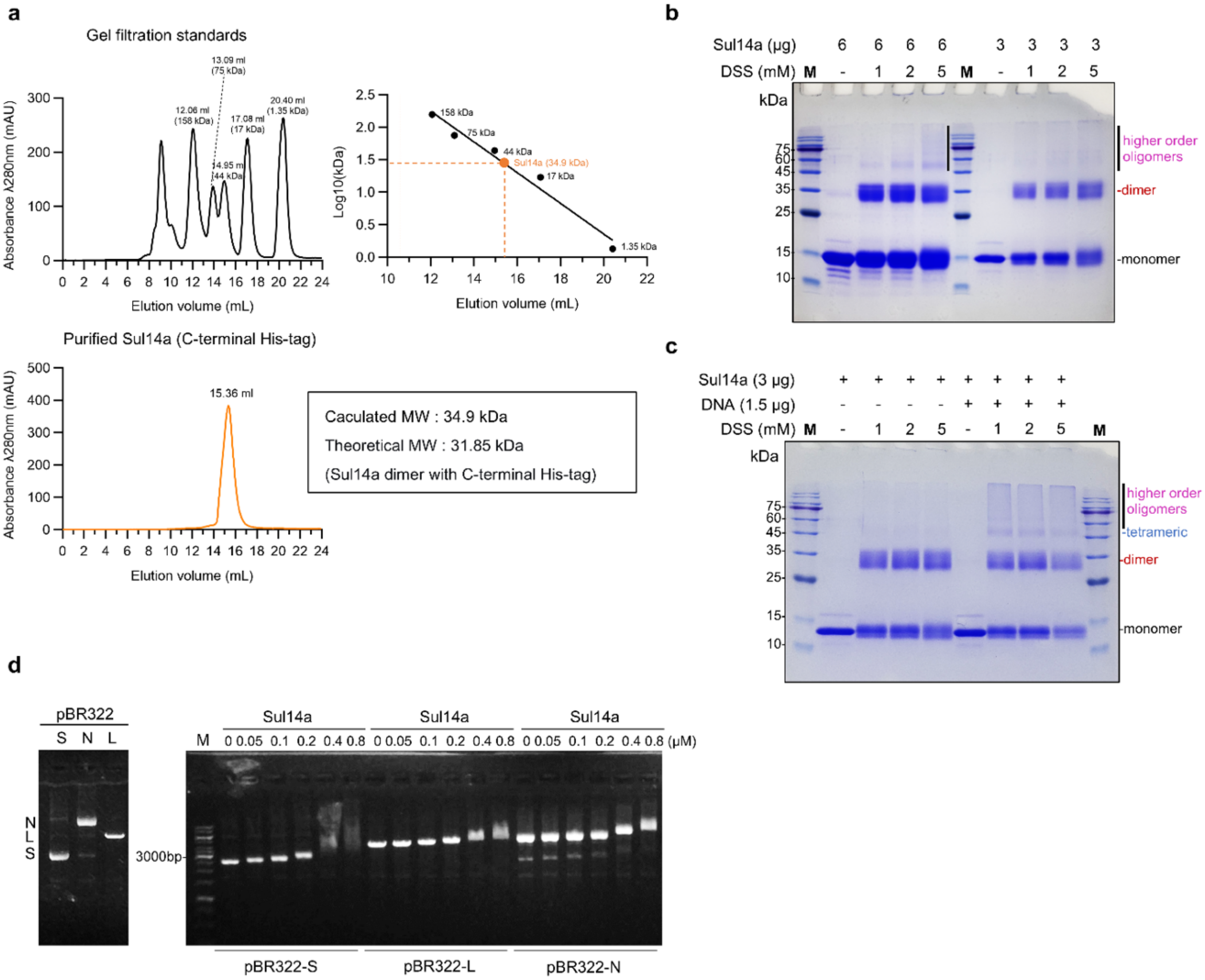


**Fig. S2. Sul14a exists as a dimer in solution, related to Fig. 2.**

**a**, Determination of the molecular size Sul14a by size-exclusion chromatography. The molecular weight of Sul14a was calculated based on its peak position (15.36 ml) in the elution profile. **b**, Chemical crosslinking of Sul14a. Purified Sul14a (3 µg and 6 µg) was incubated with increasing concentrations of DSS (1, 2, and 5 mM) in the reaction mixture, with untreated protein as a control. The samples were subjected to electrophoresis and the gel was stained. The positions of Sul14a monomer, dimer and higher order oligomers are labeled. DSS: Disuccinimidyl suberate. M: protein marker. **c**, Chemical crosslinking of the Sul14a in the presence of DNA. Different concentrations of DSS (1, 2, and 5 mM) and 3 µg Sul14a were added to the reaction mixture with or without 1.5 µg pBR322 plasmid DNA. **d,** DNA binding assay of Sul14a. Increasing concentrations of Sul14a (0, 0.05, 0.1, 0.2, 0.4, and 0.8 µM) were incubated with 300 ng DNA. Each reaction was performed with at least three technical replicates. The position of the three types of pBR322 is shown on the left panel. S: supercoiled; N: nicked; L: linearized. M: DNA marker.

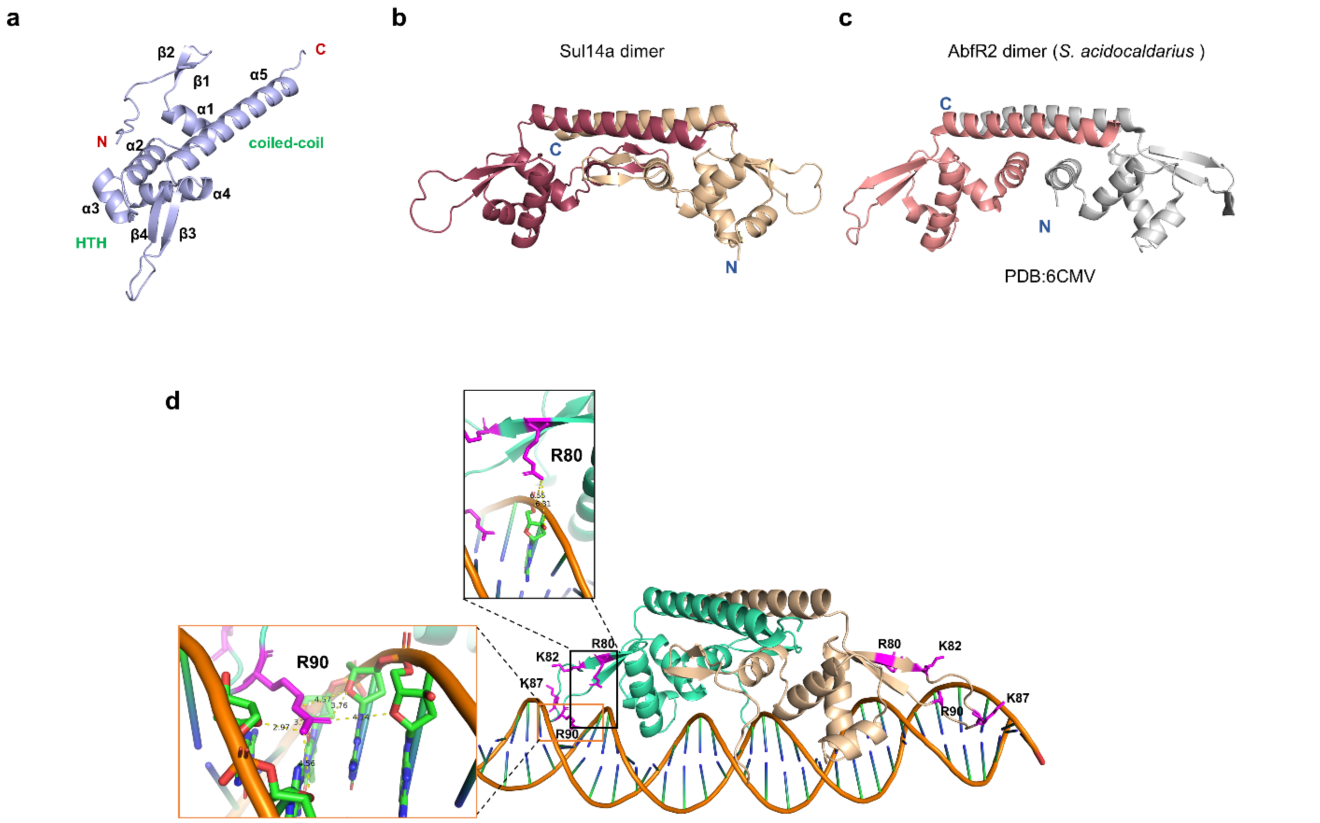


**Fig. S3. Predicted 3D structures of Sul14a and the protein (dimer)-DNA complex.**

**a,** Structure of the Sul14a monomer predicted by AlphaFold3. **b**, Structure of Sul14a dimer predicted by AlphaFold3. The brown and wheat represent Sul14a two monomers, respectively. **c**, Structure of the AbfR2 (PDB:6CMV) of *S. acidocaldarius* [1]. **d,** Structure of Sul14a (dimer)-DNA complex predicted by AlphaFold3. R80, K82, K87 and R90 are shown as sticks. Zoom in views show the predicted interactions between DNA and residues R80/R90 of Sul14a. Yellow dashed lines indicate potential hydrogen bonds and electrostatic interactions between the guanidino nitrogen atoms of the arginine residues and oxygen atoms of DNA (distances in Å). The shorter distances for R90 (2.5–4.0 Å) suggest that it has stronger interactions with DNA.

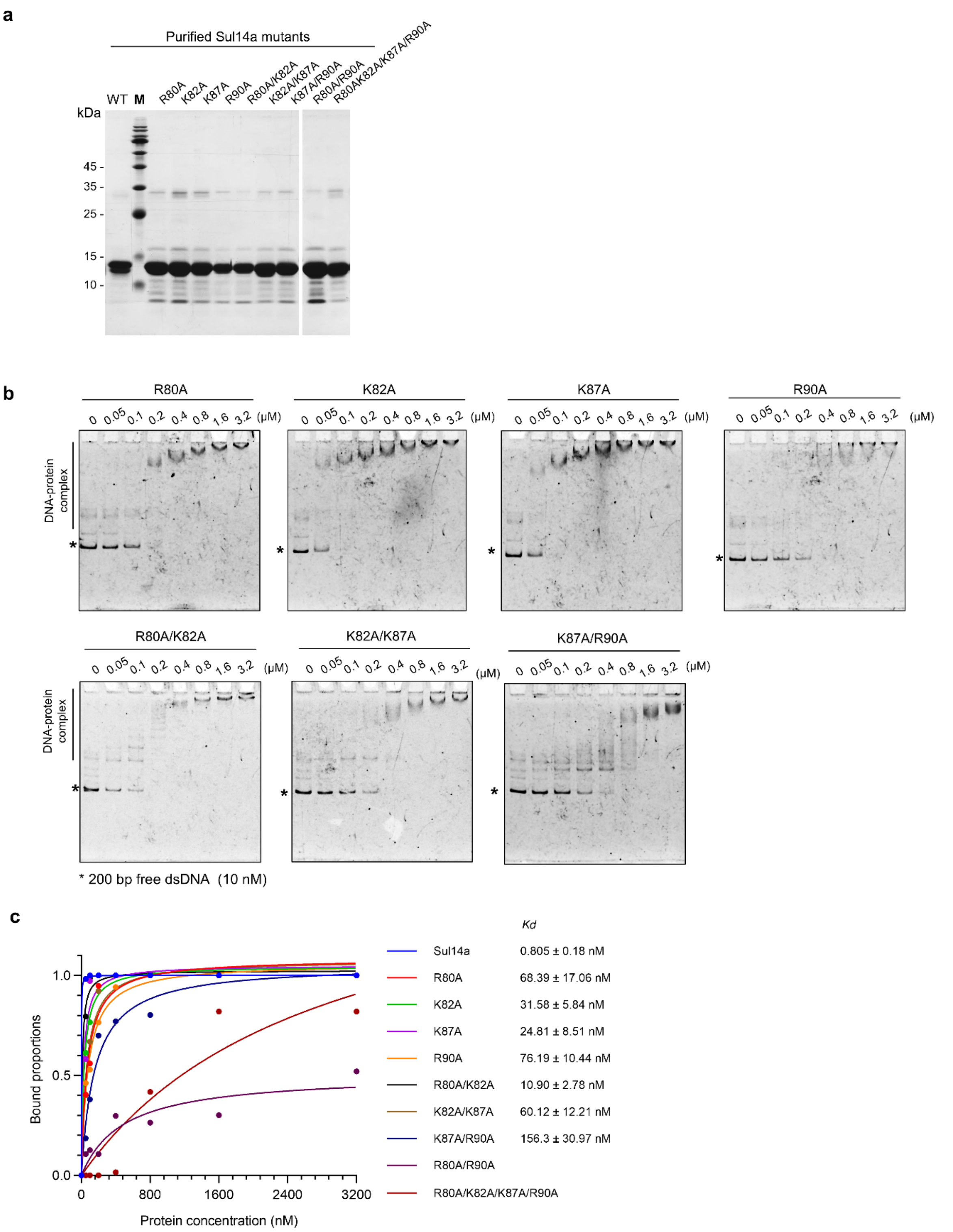


**Fig. S4. Analysis of DNA binding activity of Sul14a mutants, related to Fig. 2.**

**a**, SDS-PAGE analysis of the purified Sul14a and its mutants expressed in *E. coli*. All the protein bands matched the theoretical molecular mass of 14.3 kDa. The proteins were all fused with 6×His tag at the C-terminus. WT: wild type Sul14a. Sul14a alanine-substitution mutants are indicated. M: protein marker. **b**, EMSA analysis of the DNA binding capacity of Sul14a and its mutants. **c,** Quantitative analysis of the results in (b). The quantification was performed using ImageJ. Michaelis-Menten equation was used to fit the curve and obtain the binding constant. At least two technical replications were performed for each protein.

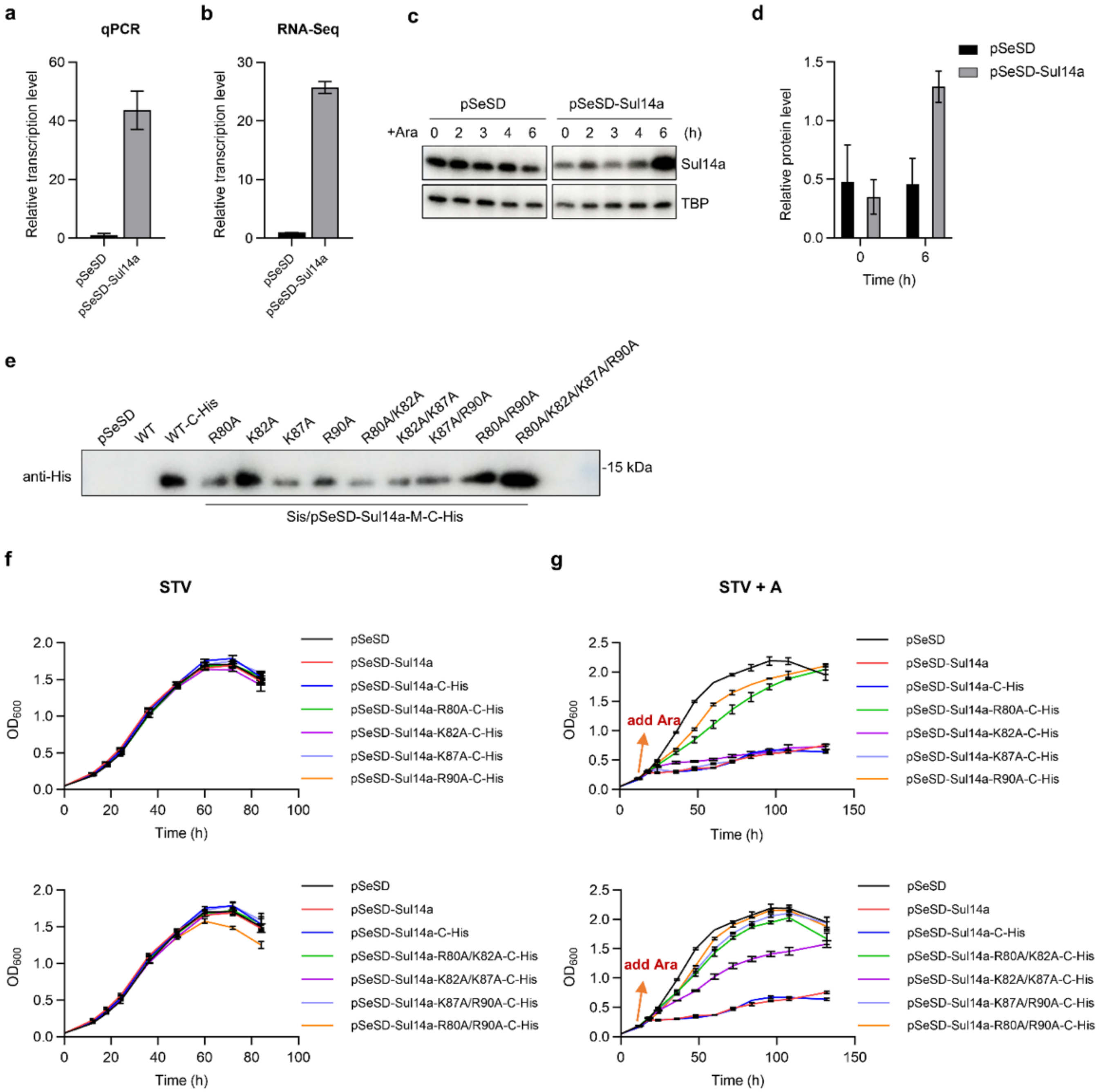


**Fig. S5. Effects of overexpression of Sul14a and its mutants on cell growth, related to Fig. 4.**

**a** and **b,** Relative mRNA level of *Sul14a* detected by qRT-PCR (a) and RNA-Seq (b) in the Sul14a overexpression strain after 6 h cultivation in arabinose-containing medium. The 16S rRNA gene was used as the internal reference. Data were normalized by the FPKM of *Sul14a* in pSeSD. The results in RNA-seq were based on three biological replicates. **c,** Western blotting analysis of the protein levels of Sul14a in the overexpression strain and control after induction. TBP (TATA-box binding protein) was used as a control. The experiments were performed at least three times, with representative images being shown. **d,** Quantification of the results in (c) by ImageJ. The relative protein levels of Sul14a at 0 h and 6 h were shown. **e**, Western blotting verification of the expression of His-tagged Sul14a mutants using anti-6×His-tag antibody. **f** and **g,** Growth curves of the mutant overexpression strains in STV (f) or ATV (g) medium. At least three biological replicates and two technical replicates were performed for each curve.

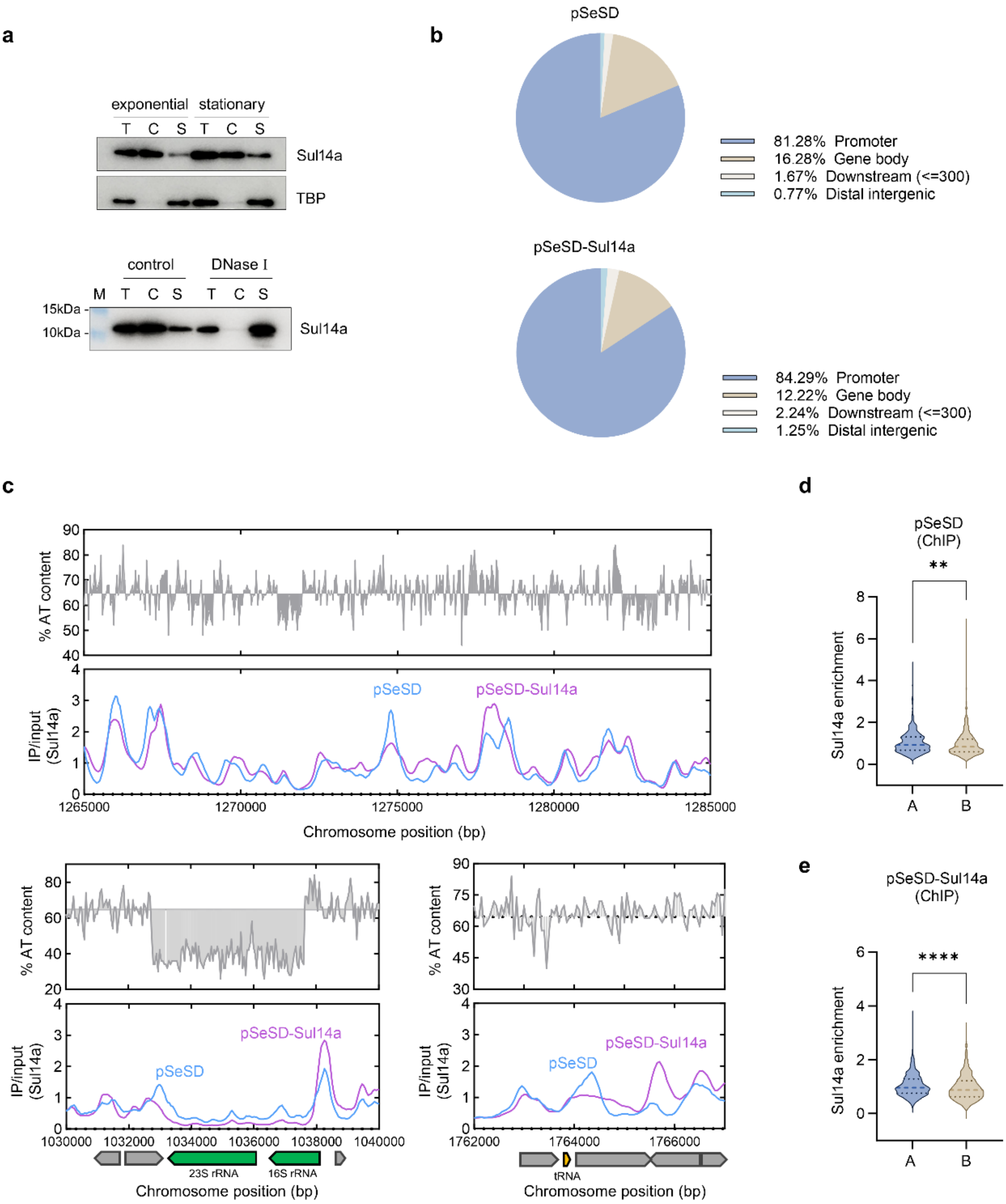


**Fig. S6. Sul14a prefers to bind to AT-rich region on the chromosome, related to Fig. 6.**

**a,** Chromatin fractionation of Sul14a in exponential and stationary phase E233S cells. Total protein (T) was separated into chromatin (C) and soluble (S) fractions, and the samples were separated by SDS-PAGE. Sul14a was detected by Western blotting using anti-Sul14a antibody. TBP was used as a soluble protein control. For samples treated with DNase I, total protein sample was incubated with DNase I before fractioning. **b**, Pie charts of the distribution of Sul14a ChIP peaks on the chromosome of cells carrying pSeSD and pSeSD-Sul14a. The majority of peaks are located in the promoter regions. **c**, Zoom in views of genomic AT content (top) and Sul14a ChIP-seq signals (bottom) in pSeSD and pSeSD-Sul14a cells. AT content below the genomic average (64.7%) is plotted in reverse. Three regions are displayed with the gene annotations being showing at the bottom of the lower panel. **d** and **e,** Analysis of the differences of Sul14a enrichment in A/B compartments between the control (d) and the overexpression strain (e). Differences were calculated using the unpaired two-tailed Student's t-test with a significance threshold of $p < 0.05$ ( $**P < 0.01$， $****P < 0.0001$).

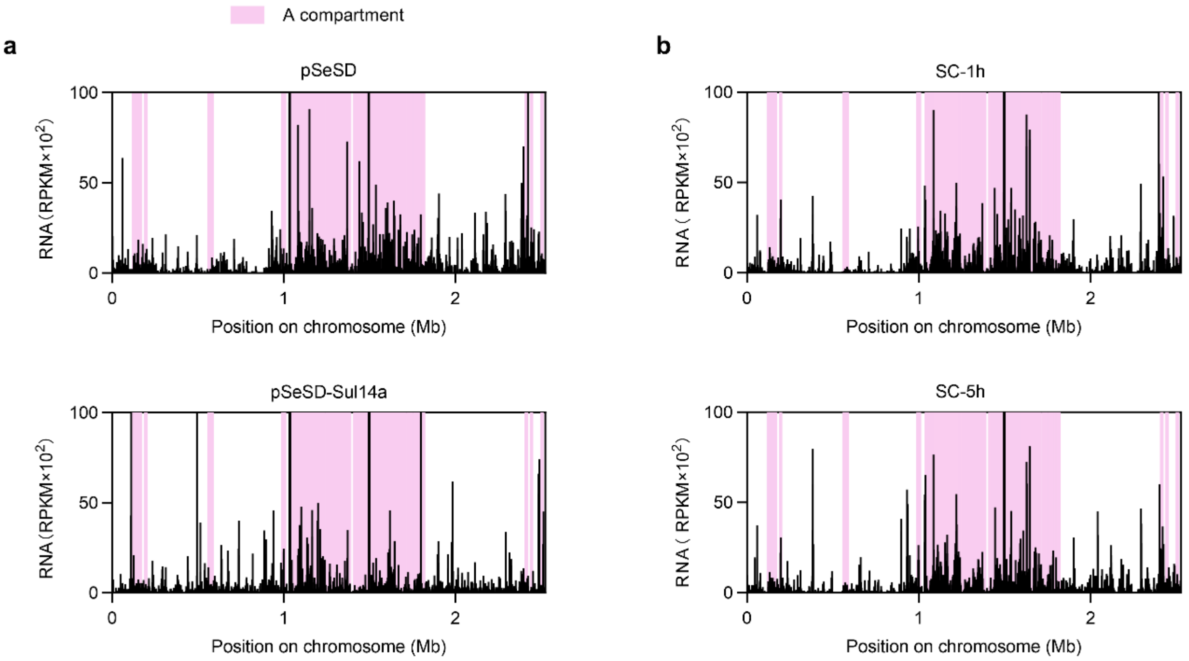


**Fig. S7. Transcript abundance in Sul14a-overexpressing and G2-synchronized cells, related to Fig 9.**

**a,** RNA-seq profiles of cells carrying pSeSD (control) and pSeSD-Sul14a. RNA transcript abundance (reads per kilobase per million mapped reads, RPKM)) is plotted for 1-kb bin. Shaded pink regions indicate the A compartment. Genome coordinate is shown on the x-axis. **b,** RNA-seq profiles of 1h and 5h cells after release from G2 synchronization. The data were derived from published sources [2].

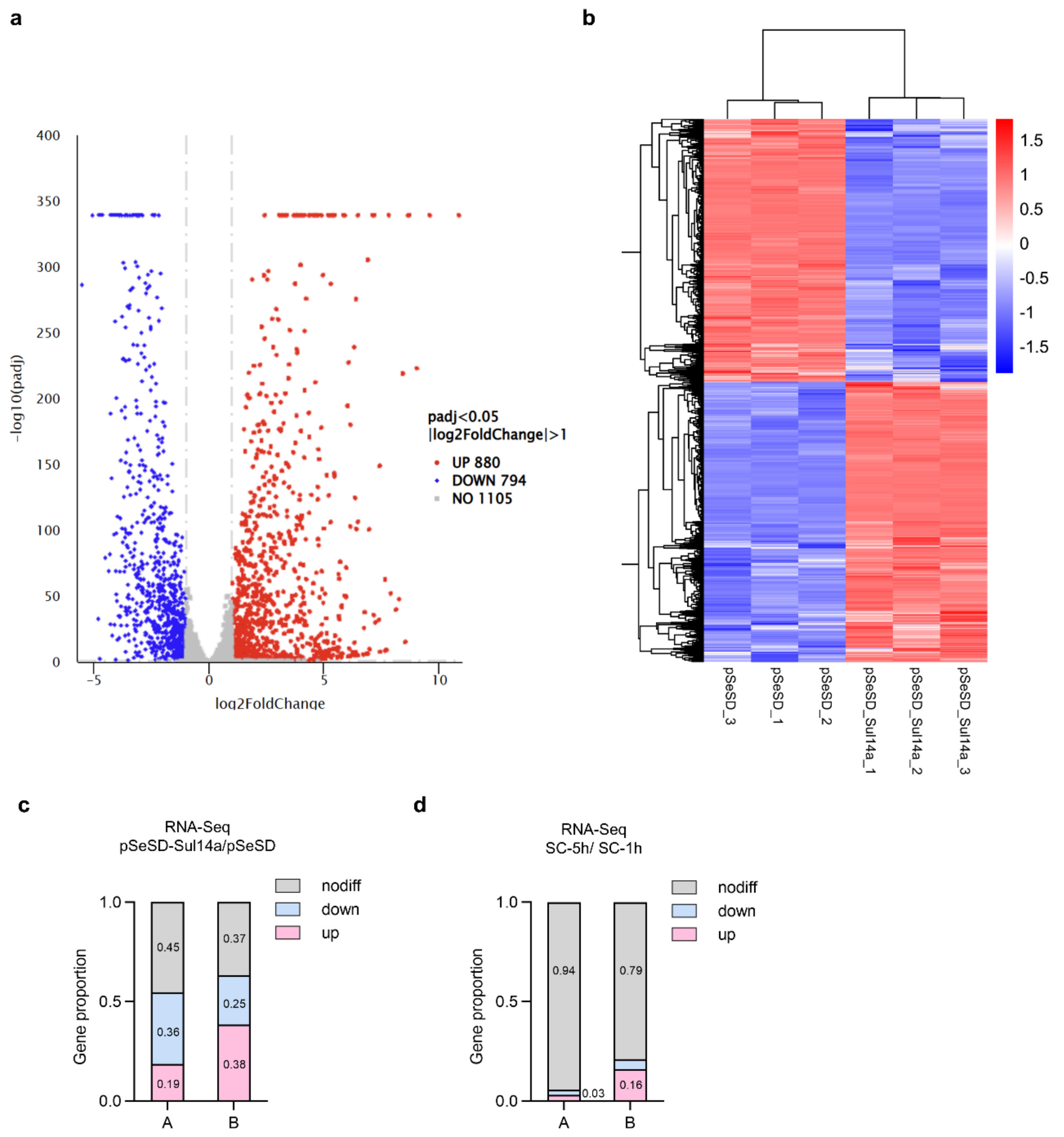


**Fig. S8. RNA-Seq analysis of Sul14a overexpression strain.**

**a,** Volcano plot showing differential gene expression in pSeSD and pSeSD-Sul14a. Both strains were grown in ATV medium for 6 h for Sul14a induction. The Y-axis (-log10, padj) represents the statistical significance of the fold change, and the x-axis represents the log2 fold change in gene expression. Genes exhibiting ≥2-fold up- and down-regulation are highlighted in red and blue, respectively. **b,** Cluster heatmap of the gene expression difference in pSeSD and pSeSD-Sul14a. Genes are clustered with log2 (FPKM + 1) and their expression levels are indicated by different colors with red representing the highest and blue indicating the lowest. **c and d,** Statistics of the global gene expression changes in A/B compartments in Sul14a overexpression cells (pSeSD-Sul14a/pSeSD) (c) and in the synchronized cells (5h/1h) (d). Changes >2 fold were regarded as notable variations. The results are based on three biological replicates.

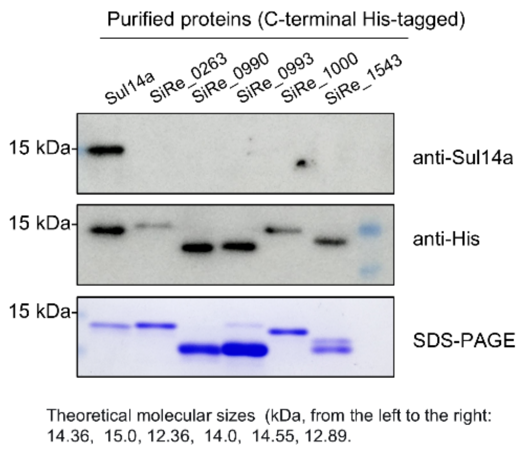


**Fig. S9. Test of the specificity of the antibody.**

Western blotting was performed for His-tagged Sul14a and its paralogs using both anti-His and anti-Sul14a antibodies.

**Supplementary Tables**

**Table S1. Lrs14-type protein homologs in representative Sulfolobales members.**

| Species / Clusters* | | *Saccharolobus islandicus* REY15A | *Saccharolobus solfataricus* P2 | *Sulfolobus acidocaldarius* DSM639 | *Sulfurisphaera tokodaii* str. 7 | *Metallosphaera sedula* DSM5348 |
|---|---|---|---|---|---|---|
| Cluster 1 | | SiRe_1543 | SSO0458 (Smj12) | Saci_0446 (AbfR1) | ST0173 | Msed_2175 |
| Cluster 2 | a | SiRe_1000 | SSO1101 | Saci_1219 | ST0837 | Msed_1397 |
| | b | SiRe_0263 | SSO2474 | Saci_1223 (AbfR2) | ST0749 | - |
| | c | - | - | - | ST1043 | - |
| Cluster 3 | | SiRe_1948 (Sul14a) [#] | SSO0048 (Sta1) | Saci_0102 | ST2050 | Msed_2027 |
| Cluster 4 | a | SiRe_0993 | SSO1110 | Saci_0133 | ST1867 | Msed_1566 |
| | b | - | - | - | ST1890 | - |
| Cluster 5 | a | SiRe_0990 | SSO1108 (Lrs14) | - | ST0980 | Msed_1563 |
| | b | - | - | Saci_1242 | ST1889 (Sto12a) | - |

* The five Lrs14 clusters marked in the table are according to Veerke De Kock et al (2025) [3].

### The protein investigated in this study.

**Table S2. Strains and plasmids used in this study**

| Strain or vector | Description | References or sources |
|---|---|---|
| Escherichia coli DH5α | Plasmid amplification | Laboratory strain |
| E. coli BL21(DE3) codon plus-RIL | Protein expression | Laboratory strain |
| Sa. islandicus REY15A (E233S) | ΔpyrEFΔlacS | Deng et al.[4] |
| pSeSD | A *Saccharolobus-E. coli* shuttle vector carrying an expression cassette controlled under a synthetic strong promoter $P_{araS\text{-SD}}$ | Peng et al.[5] |
| pSeSD-Sul14a | pSeSD carrying wild-type Sul14a encoding sequence | This study |
| pSeSD-Sul14a-C-His | Expression of Sul14a with His-tag at the C-terminal | This study |
| pSeSD-Sul14a-R80A-C-His | pSeSD carrying Sul14a encoding sequence with R80A substitution | This study |
| pSeSD-Sul14a-K82A-C-His | pSeSD carrying Sul14a encoding sequence with K82A substitution | This study |
| pSeSD-Sul14a-K87A-C-His | pSeSD carrying Sul14a encoding sequence with K87A substitution | This study |
| pSeSD-Sul14a-R90A-C-His | pSeSD carrying Sul14a encoding sequence with R90A substitution | This study |
| pSeSD-Sul14a-K82A/K87A-C-His | pSeSD carrying Sul14a encoding sequence with K82A/K87A substitutions | This study |
| pSeSD-Sul14a-R80A/K82A-C-His | pSeSD carrying Sul14a encoding sequence with R80A/K82A substitutions | This study |
| pSeSD-Sul14a-K87A/R90A-C-His | pSeSD carrying Sul14a encoding sequence with K87A/R90A substitutions | This study |
| pSeSD-Sul14a-R80A/R90A-C-His | pSeSD carrying Sul14a encoding sequence with R80A/R90A substitutions | This study |
| pSeSD-Sul14a-R80A/K82A/K87A/R90A-C-His | pSeSD carrying Sul14a encoding sequence with R80A/K82A/K87A/R90A substitutions | This study |
| pET22b-Sul14a-K82A-C-His | pET22b carrying Sul14a encoding sequence with K82A substitution | This study |
| pET22b-Sul14a-K87A-C-His | pET22b carrying Sul14a encoding sequence with K87A substitution | This study |
| pET22b-Sul14a-R90A-C-His | pET22b carrying Sul14a encoding sequence with R90A substitution | This study |
| pET22b-Sul14a-K82A/K87A-C-His | pET22b carrying Sul14a encoding sequence with K82A/K87A substitutions | This study |

| | | |
|---|---|---|
| pET22b-Sul14a-R80A/K82A-C-His | pET22b carrying Sul14a encoding sequence with R80A/K82A substitutions | This study |
| pET22b-Sul14a-K87A/R90A-C-His | pET22b carrying Sul14a encoding sequence with K87A/R90A substitutions | This study |
| pET22b-Sul14a- R80A/R90A-C-His | pET22b carrying Sul14a encoding sequence with R80A/R90A substitutions | This study |
| pET22b-Sul14a-R80A/K82A/K87A/R90A-C-His | pET22b carrying Sul14a encoding sequence with R80A/K82A/K87A/R90A substitutions | This study |

**Table S3. Oligonucleotides used in this study**

| Primers | Sequence[a,b] (5'-3') |
|---|---|
| Sul14a-NdeI-F | GTGGAATTCATATGATGTCGGAAACCCAATT |
| Sul14a-SalI-R | AGGCGTCGATTACAATGGCTTAAACTCCTTT |
| Sul14a-SalI-C-His-R | AGGCGTCGACAATGGCTTAAACTCCTTT |
| *Sul14a*-R80A/K82A/K87A/R90A-SOE-F | ATTAGTAATG**GCA**ATA**GCG**GAGCCAGGGAAC**GCG**GCTGGA**GCA**CCAAGATATTTA |
| *Sul14a*-R80A/K82A/K87A/R90A-SOE-R | ATAAATATCTTGG**TGC**TCCAGC**CGC**GTTCCCTGGCTC**CGC**TAT**TGC**CATTACTAAT |
| *Sul14a*-R80A/R90A-SOE-F | ATTAGTAATG**GCA**ATAAAGGAGCCAGGGAACAAGGCTGGA**GCA**CCAAGATATTTA |
| *Sul14a*-R80A/R90A-SOE-R | ATAAATATCTTGG**TGC**TCCAGCCTTGTTCCCTGGCTCCTTTAT**TGC**CATTACTAAT |
| *Sul14a*-K82A-SOE-F | ATTAGTAATGAGAATA**GCG**GAGCCAGGGAACAAGG |
| *Sul14a*-K82A-SOE-R | CCTTGTTCCCTGGCTC**CGC**TATTCTCATTACTAAT |
| *Sul14a*-K87A-SOE-F | GGAGCCAGGGAAC**GCG**GCTGGAAGACCAAGATA |
| *Sul14a*-K87A-SOE-R | TATCTTGGTCTTCCAGC**CGC**GTTCCCTGGCTCC |
| *Sul14a*-R80A-SOE-F | GATGGGATTAGTAATG**GCA**ATAAAGGAGCCAGGGA |
| *Sul14a*-R80A-SOE-R | TCCCTGGCTCCTTTAT**TGC**CATTACTAATCCCATC |
| *Sul14a*-R90A-SOE-F | AGGGAACAAGGCTGGA**GCA**CCAAGATATTTATAC |
| *Sul14a*-R90A-SOE-R | GTATAAATATCTTGG**TGC**TCCAGCCTTGTTCCCT |
| qPCR- *Sul14a*-F | CCTTGTTCCCTGGCTCCTTT |
| qPCR- *Sul14a*-R | TGAAAGGCGATGCTAGAGGT |
| qPCR-16S-F | CGCAAGACTGAAACTTAAAGGA |
| qPCR-16S-R | AGTCAGGCAAGGTCGTTAG |
| Biotin-dbr-F | ATCTGTAATTGCCTTAAGCCCAACTTTA |
| dbr-1000bp-R | ATTGCCTCAACCAAAAGGAAACTCCCCA |
| FAM-AT-F | TTAATATTATTTAATATTTTATAAAATTTTTATTAATTTAATTTTTAATTATTTT |
| AT-R | AAAATAATTAAAAATTAAATTAATAAAAATTTTATAAAATATTAAATAATATTAA |
| FAM-GC-F | GCGGCCGGGCCCGGGCCCCGCCGGGCCCCGCCGGGCCCCGCCGGGCCCCGCGCCG |
| GC-R | CGGCGCGGGGCCCGGCGGGGCCCGGCGGGGCCCGGCGGGGCCCGGGCCCGGCCGC |

[a]The underlined denote sites of the restriction enzymes. [b]The mutated codons are indicated in boldface

#### Supplementary Materials and Methods

##### Bioinformatic analysis

Homology searches were performed using Protein BLAST (National Center for Biotechnology Information). Clustal Omega version 1.2.4 and ESPript 3.0 was used to generate sequence alignment [6, 7]. The phylogenetic tree was generated by Maximum Likelihood method with the LG+G model by IQ-Tree 2 [8]. SWISS-MODEL [9] and AlphaFold 3.0 [10] were used for protein structure prediction.

##### Generation of the overexpression strains of Sul14a mutants

*Sul14a* and mutant derivatives were engineered by PCR to have a C-terminal 6×His-tag sequence. The *Sul14a* gene and its mutant fragments were cloned into pSeSD plasmids digested by *NdeI* and *SalI*. The plasmids were transformed into E233S by electroporation, generating strains named pSeSD-Sul14a-R80A-C-His, pSeSD-Sul14a-R90A-C-His, pSeSD-Sul14a-K82A-C-His, pSeSD-Sul14a-K87A-C-His, pSeSD-Sul14a-R80A/K82A-C-His, pSeSD-Sul14a-K82A/K87A-C-His, pSeSD-Sul14a-K87A/R90A-C-His and pSeSD-Sul14a-R80A/R90A-C-His for short. Growth curves of Sul14a mutant overexpression strains were obtained by inoculating cells in STV or ATV medium from an initial $OD_{600}$ of 0.03-0.05 and subsequent culture and measurement. The samples were taken every 6 or 12 h to get the $OD_{600}$ values. Cells containing pSeSD were used as the control.

##### Electrophoretic mobility shift assay (EMSA)

Circular supercoiled pBR322 plasmids were purified using a Plasmid Extraction Kit (Omega Bio-tek, Norcross, USA) following the provided protocol with two additional wash steps. linearized and nicked pBR322 plasmid DNAs were prepared by treatment with *BamHI* (Takara Biomedical Technology, Beijing) and Nb. *Bpu10I* (Thermo Fisher Scientific), respectively, and were purified by using a DNA Cycle-Pure Kit (Omega Bio-tek, Norcross, USA). For the reaction with plasmid DNA as substrate, different concentrations of Sul14a (0, 0.05, 0.1, 0.2, 0.4, and 0.8 μM) were taken with 300 ng of plasmid pBR322 with different types (supercoiled, linearized, and nicked), and the reaction system was 50 mM Tris-HCl pH 6.8, 25 mM NaCl. The reaction was incubated at room temperature or 37°C for 30 min. After incubation, the results were detected by 1% agarose gel electrophoresis with 0.5×TAE buffer at 110 V, 40 min, and visualized by EC3 Imaging System (Ultra-Violet Products Ltd, Cambridge UK).

##### Protein chemical cross-linking

A quantity of 3 μg or 6 μg Sul14a was cross-linked with DSS (Disuccinimidyl suberate) crosslinker (Thermo Scientific) with a final concentration of 0, 1, 2, and 5 mM in solution.Reactions including DNA were performed with the addition of 1.5 μg of pBR322 DNA and 3 μg of Sul14a protein. The mixtures were incubated at room temperature for 30 min. Then the samples were treated by adding 5×loading buffer (with 2-Mercaptoethanol) and boiling for 10 min before fractionating by 15 % sodium dodecyl sulfate-polyacrylamide gel electrophoresis (SDS-PAGE).

##### Chromatin fractionation

The chromatin fractionation assay was performed as described [11]. Two portions of *Sa. islandicus* E233S cells with a total $OD_{600}$ of about 2 were collected separately at $OD_{600}$ = 0.2 for exponential samples and $OD_{600}$ = 1.0 for stationary samples. The samples were resuspended with chromatin extraction buffer (25 mM HEPES pH 7.5, 15 mM $MgCl_2$, 100 mM NaCl, 400 mM Sorbitol, 0.5% Triton X-100) and incubated on ice for 10 minutes. The lysates were centrifuged (14,000 × *g*, 20 min, 4°C) to separate soluble (supernatant) and chromatin (pellet) fractions. Chromatin pellets were resuspended in equal-volume buffer and sonicated 10 times on ice with a JY92-II sonicator (Scientz Biotechnology, Ningbo, China). The two fractions and the whole cell extracts were resolved by SDS-PAGE and immunoblotted with Sul14a antibody. In addition, set up a control group treated with DNase I for the exponential cells.

**The DNA sequence information**

55bp AT dsDNA for EMSA:

TTAATATTATTTAATATTTTATAAAATTTTTATTAATTTAATTTTTAATTATTTT

55bp GC dsDNA for EMSA:

GCGGCCGGGCCCGGGCCCCGCCGGGCCCCGCCGGGCCCCGCCGGGCCCCGCGCCG

200bp 5’-FAM-dsDNA for EMSA in Fig.2 and Fig.S3:

>Promoter of *sire_0124*

GTAAATAATATAATATGTAATATTATCATATATAACCTTGTATTCTAGAAAGACGGGAGTCATTAAACATTACTTGG
AAACAGTTACTTAAAAAATAGCCTAAAGTCATGTAGACTGGTAGTCGAAAATCAGAGAGAGGTTTTTAAAAGT
GTTCTGAGAAACGGATTTCAGCAAGTGATCTAAAAATGAACACCAAAAAA

500bp 5’-FAM-dsDNA for DNA bridging assay in Fig.4 and for EMSA in Fig.S8:

>500bp from gene *sire_1550*

TCGTATTCCGGATCTTCATCTTCTTTAATGAACGTGTTTGGCATATTACTTATTTTAGATTCTAATAATTTAAACTCT
TCACATAATGTGTTTGATCTTATAAGAGTTAATGAGGTAAGTTATATATTAGCCATGCTCGGAAAAGATTTCCAAA
AGATTTGGGGTTATGAGCAGGAGAGATTTAAAATAAAGGGTTCTAAGGAGCCTTTAAAGTATAGGTTAGTTAAC
GCTCACTATAAAATTAGCTCCATGATTAGTAGGCTTGACGCTTACATTTCTAGAATGCAAGAAAGGGATAAAATA
CTATTTGAGAGAGTAGTAGAAGCACAAATGTCTAAGGATACACAAAGGGCAGCAATGTACGCTAATGAGGTAG
CAGAAATCAGAAAGATCTCCAGACAGTTGTTAACAACCCAAATTGCCTTGGAACAAGTACAACTCAGATTAGA
GACAGTAACTGAATTAGGAGATGTATTTGTAAACTTAATCCCAGTATTAGG

1000bp biotin-labeled-dsDNA (amplified by PCR):

ATCTGTAATTGCCTTAAGCCCAACTTTATCGTCTATCATATTTACAGTGATATTATATTTTAGAATTATTTCCAAAAT
TTGTACCGAATACTCTAAATTACTAGTTATTTTAAGGATTCTCCCTGTTAATTTCACTACGAAAGGATTCATATATG
AATATAGTTTTTTCGAGTATTATGCCTTATCGTACTTAAAAAGCGTAGACTATCAAAGATGTAGATTAGAGCCGAA
ATGGCCTCAAGTAAAGTGGAAGATTTCGTAAAGAATTGGGGAGGTAAACAAGAGCCAAGTATTGGTGAAAGA
ATAAAGAATGCGTTCAAGCCACAGCAACCACTAAGGTATAGATTAGTAATGGCAAACTACAGATTAAGGACAAT
GGTAAGTCGTCTTGACGTTTACATTTCAAGATTACAAGAGAGAGATAGGACTCTATTCGAAAAGGTCGTAGAAT
CTCAGATGTCAAAGGATACGGCAAGAGCAGCTATGTATGCTAATGAAATAGCTGAGATTAGGAAGATTTCCAGG
CAGCTAATTACTACACAAATTGCCCTAGAGCAAGTACAACTCAGGTTAGAGACGATAACTGAGCTTGGAGATGT
ATTTAACAGCCTAATACCAGTACTTGGTGTTATAAAAGAGCTAAGAAATGCGATGAAAGGAGTTATGCCAGAGA
TAAGTTTGGAGCTAGCAGAATTAGAGGAGGGATTACAAGAGGTAGTAATAGAAGCAGGGGACTTTACTGGTGC
GCCAGCCAACTATGGTGCTTCAAGCCCAGAGGCAAGGAAGATATTAGAAGAAGCTTCTGTTGTTGCTGAGCAG
AGAATGAAAGAGAAGTTCCCAGAATTGCCAAGCTTCGTTACCTCTACTCAGAAAGTATCTAATCAAGAGCAGA
AATAATATCTGATTCATAGATTTTTTCCTAAGATTTATACGGTAAAATAGTTTTTTAAGGTTTGGATTGGTAATATAT
ATTGGGGAGTTTCCTTTTGGTTGAGGCAAT

**Legends for the supplementary data**

Supplementary data 1: Dataset of the ChIP_Seq enrichment peaks of cells carrying pSeSD and pSeSD-Sul14a.

Supplementary data 2: Dataset of the RNA_seq analysis of cells carrying pSeSD and pSeSD-Sul14a.

Supplementary data 3: Source data used to generate the graphs in this study.

Supplementary data 4: Uncropped blots/gel images with corresponding molecular size marker.
